# Linker Histones Enhance Robustness in Diurnal Transcription Dynamics

**DOI:** 10.1101/2025.06.26.661703

**Authors:** Kinga Rutowicz, Lena Perseus, Marc W. Schmid, Jasmin Schubert, Diana Zörner-Pazmino, Tomasz Bieluszewski, Maciej Kotlinski, Ricardo Randall, Piotr Ziolkowski, Andrzej Jerzmanowski, Clara Bourbousse, Fredy Barneche, Célia Baroux

## Abstract

Linker histone H1 is crucial for chromatin organization and gene expression in *Arabidopsis thaliana*, influencing development and stress responses. To explore its role in diurnal gene regulation, we examined H1-deficient plants and found that H1 is essential for maintaining rhythmic gene expression. Genes losing synchronization often contained NAC transcription factor binding sites, indicating H1 may affect their accessibility. Nuclear imaging revealed that H1 subtly modulates nuclear size and chromatin distribution across the photoperiod. Epigenetic analysis showed typical diurnal changes—declines in H3K4me3 and active RNA Pol II in the evening and increases in H3K27me3. In H1 mutants, these patterns persisted but with elevated H3K4me3 and RNA Pol II (Ser2P) levels at night and in the morning. These results suggest that H1 fine-tunes chromatin and transcriptional rhythms, contributing to the temporal coordination of gene activity in response to environmental and developmental signals.

## Introduction

Diurnal adaptation in plants involves a coordinated set of responses to optimize growth and resource allocation across the day-night cycle, as well as the vegetative-to-reproductive transition (Coen & Prusinkiewicz, 2024; Nozue & Maloof, 2006; Venkat & Muneer, 2022; Wang et al., 2024). At the physiological level, plants adjust key processes such as photosynthesis, respiration, and stomatal movement, water and ion transport (Matthews et al., 2018; Venkat & Muneer, 2022). At the molecular level, diurnal variations are detected in the proteome, the transcriptome and the epigenome (Nozue & Maloof, 2006; Wang et al., 2024) and references below). While some of these rhythms are regulated by circadian rhythms and remain stable despite temporary disturbances, others respond directly to environmental cues that vary throughout the day-night (diurnal) cycle (Nozue & Maloof, 2006).

In Arabidopsis, diurnal variations of the proteome influence a variety of processes including light perception, photosynthesis, metabolism and ion transport (Uhrig et al., 2021). Diurnal variations extend to protein isoforms, such as acetylated and phosphorylated forms, which are crucial for the proteins’ functional activity (Uhrig et al., 2021; Uhrig et al., 2019). Protein translation, which, along with protein degradation, plays a critical role in controlling the abundance of proteins available to the cell, also shows diurnal variation (Duncan & Millar, 2022; Li et al., 2022; Seaton et al., 2018; Uhrig et al., 2021). Protein abundance also correlates with transcriptional fluctuations, a process called translational coincidence (Seaton et al., 2018). Moreover, diurnal transcriptome dynamics also include the occurrence of different transcript isoforms because of diurnal variation in splicing activity controlled by the circadian clock (Yang et al., 2020). Further, the pattern of diurnal transcriptional changes varies with age and development suggesting a cross-talk between environmental signals and intrinsic cues on the physiological state of the plant (Jung et al., 2024) (Redmond et al., 2024).

In mice, transcriptional oscillations can be largely explained by rhythmic RNA Polymerase II (RNA Pol II) recruitment at promoters rather than rhythmic transition from paused to productive transcript elongation (Le Martelot et al., 2012). In plants, RNA Pol II also exhibits genome-wide rhythmic occupancy, also shown in Rice to correlate with changes in the spatial organization of the genome (Deng et al., 2022). Core clock genes show high spatial proximity in the nuclear space (chromatin connectivity) in the morning and weaker connectivity in the evening, indicating a shift from coordinated transcription in the morning to discretely regulated transcriptional regions in the evening (Deng et al., 2022). Likewise, chromatin accessibility differs in Arabidopsis in response to photoperiod (Tian et al., 2021). Such diurnal changes in chromatin organization are also reflected at the epigenetic level as shown by studies conducted on *Arabidopsis thaliana* and *Brassica napus* (Baerenfaller et al., 2016; Deng et al., 2022; Fung-Uceda et al., 2024; Maric & Mas, 2020; Perales & Mas, 2007; Song et al., 2019). Notably, morning-phased or evening-phased genes show concomitant changes in the accumulation of transcriptionally permissive marks such as H3K4me3, H3K9ac, H3K27ac and H3S28p (Baerenfaller et al., 2016; Song et al., 2019). Among those genes, are transcription factors, circadian regulators and components of the starch catabolic pathway (Baerenfaller et al., 2016). By contrast, diurnal fluctuations in H3K27me1 or DNA methylation, associated with a transcriptionally repressive state, affect a different subset of genes, including cell cycle and DNA damage response genes (Balcerowicz, 2024; Fung-Uceda et al., 2024).

Spatial reorganization of the genome, changes in chromatin accessibility and epigenetic composition along the diurnal rhythm are likely involving general chromatin architects to either enable smooth transitions, stabilize new states, or both. Here, we explored the role of H1 linker histones (H1) in diurnal transcriptome fluctuations and discovered that they are crucial for coordinating the expression of genes with diurnal rhythms. Notably, genes that lose diurnal synchronization in the absence of H1 display binding sites for NAC transcription factors (TF), suggesting that H1 may influence their regulation by modulating accessibility to these TFs. Using large-scale quantitative imaging of nuclei isolated at different time points during the photoperiod, we observed subtle variations in nuclear size and chromatin distribution that were influenced by H1. We also found a decrease in global H3K4me3 and active RNA Pol II in the evening, accompanied by an increase in H3K27me3, with a role for H1 in H3K27me3 accumulation in the evening and RNA Pol II (Ser2P) in the late night and morning in the mutant.

Collectively, this study revealed a role for linker histone (H1) in regulating diurnal transcriptome and chromatin variations in Arabidopsis, focusing on how they might influence the coordinated expression of genes with rhythmic patterns. These findings pave the way to investigate the role of general chromatin architects on transcription and chromatin organization oscillations.

## Results

### H1 depletion affects transcriptional diurnal dynamics

The Arabidopsis genome encodes three H1 variants: H1.1, H1.2 and H1.3, the latter being stress-inducible and expressed only in guard cells under regular growth conditions (Wierzbicki & Jerzmanowski, 2005). Loss-of-function mutant lines were first obtained by expressing long interference RNAs (Wierzbicki & Jerzmanowski, 2005). Thereafter, T-DNA insertion lines were isolated and combined to create a *3h1* - triple homozygous mutant line (Rutowicz et al., 2019). Because this line was created by introgressing mutant alleles from different accessions (L*er* and Col_*0*, respectively), we generated a new mutant line in a Col_0 background using CRISPR-Cas9 gene editing (Bieluszewski et al., 2022). *3h1*^*crispr*^ harbours a small deletion in each of the three *H1* genes leading to frameshift mutations (**Figure 1A, Supplemental File 1**). Efficient depletion of the whole H1 complement was confirmed by the loss of detectable H1 in immunostained *3h1*^*crisp*^ nuclei (**Figure 1B**). The *3h1*^*crispr*^ knock-out also expectedly reproduces the loss of conspicuous chromocenters in which heterochromatin is compacted, early flowering phenotype under long days and increased lateral root number described previously (Rutowicz et al., 2019) (**Figure 1B, Supplemental File 1**, and below). Consequently, all experiments in this study were conducted using this new mutant line, thereafter, referred to as *3h1*.

**Figure 1.**
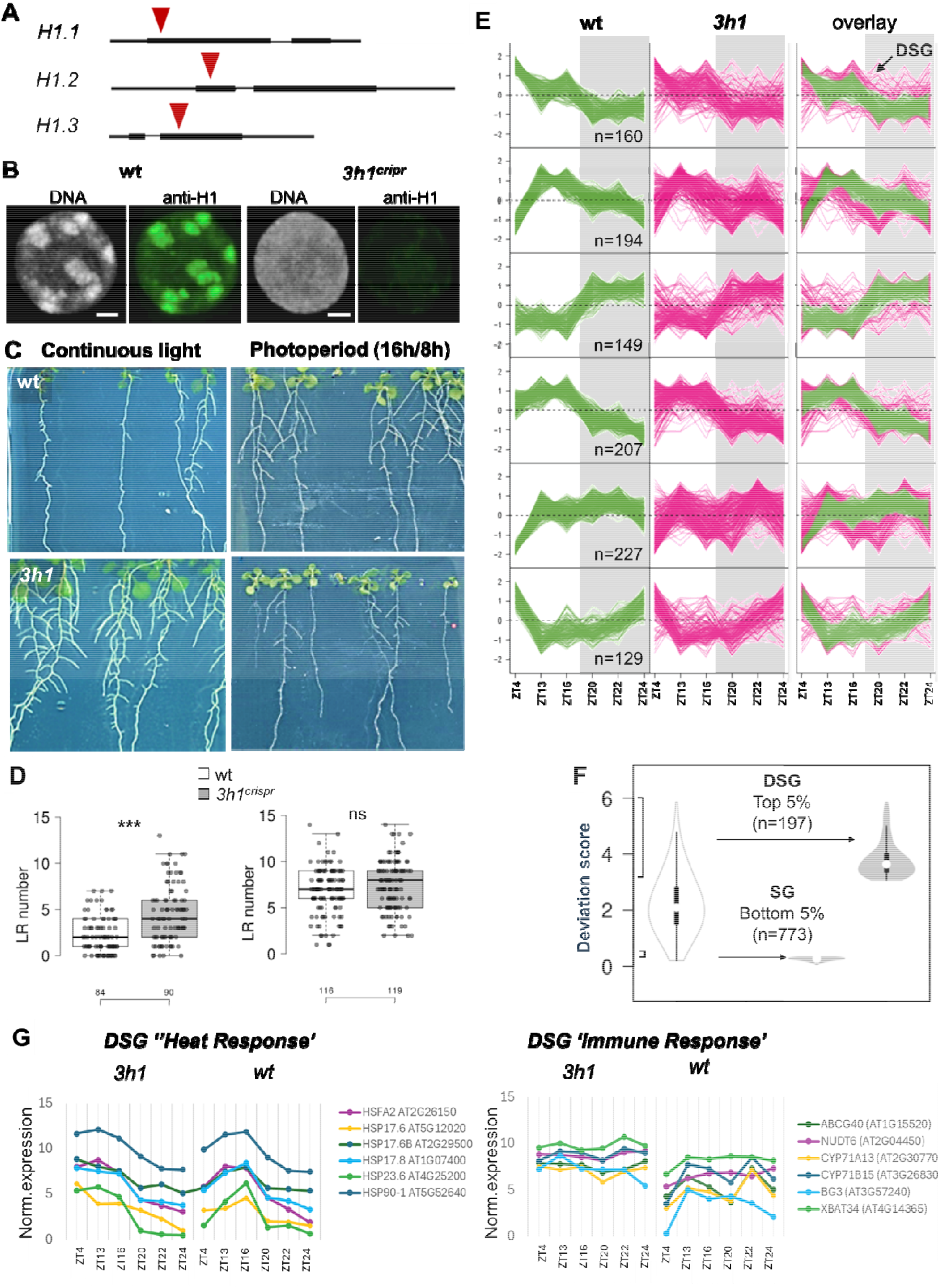
H1 depletion affects the diurnal control of gene expression. **(A)** Schematic representation of the CRISPR Cas9-mediated deletions in the *H1*.*1, H1*.*2* and *H1*.*3* coding regions. H1 Immunostaining (green) and DNA counterstaining (grey) of *3h1*^*crispr*^ leaf nuclei confirm H1 depletion and heterochromatin decondensation, respectively, as previously shown for the T-DNA-based *3h1* mutant (Rutowicz et al, 2019). Scale bar: 2µm **(C)** Representative images of 12 days-old wt and *3h1* mutant seedlings grown under continuous light or under long day conditions as indicated. (**D**) Number of lateral roots per seedling (dot) grown under continuous light (7 days old seedlings, left) or under long day photoperiod conditions (12 days old seedlings, right), for wild-type segregant seedlings (white) and *3h1 crispr* mutant seedlings (grey). ***, P<0.001, Kruskal-Wallis test with Bonferroni correction. The numbers of seedlings scored are indicated on the X axis. **(E)** Genes with co-regulated expression in wild-type (wt) seedlings were partitioned in 6 clusters (n, number of genes in each cluster). The graphs show their expression in 19dag seedlings, wt (green traces) and mutant (*3h1*, pink traces), sampled at six time points as indicated. The overlay (right) allows visualising genes losing synchronised, co-expression in the mutant (DSG, De-Synchronised Genes) in contrast to the bundled traces corresponding to genes with synchronised co-expression (SG, Synchronised Genes). **(F)** Deviation score -measuring the cumulative differences in expression of diurnally regulated genes in mutant seedlings across all time points – for all genes (left) and for the top and bottom 5% defining the DSG and SG classes, respectively. **(G)** Example of DSG expression in 3h1 and wt of the functional categories “response to heat (GO:0009408)” and “regulation of innate immune response (GO:0045088)”. The values correspond to averaged, normalised expression levels. See also **Supplementary Data 1** and **Supplementary Figure S1**.

Under continuous light, the 3h1 mutant displays an increased number of lateral roots, while overall root length remains comparable to the wild type (**Supplementary Figure S1A**). However, this excess of lateral roots is no longer apparent under a long-day photoperiod (**Figure 1C-D, Supplementary Figure S1A, B**). Notably, when both lateral root primordia and elongated lateral roots were scored, 3h1 still showed a higher initiation rate of lateral roots (**Supplementary Figure S1B**). These findings suggest that although *3h1* retains the capacity to initiate more lateral roots, their subsequent growth may be impaired under long-day conditions compared to under continuous light. The contrasting root behaviour—enhanced lateral root growth under continuous light but reduced growth under a long-day photoperiod—led us to hypothesize that H1 may play a broader role in regulating diurnal transcriptional dynamics. To test this, we analyzed the transcriptome of wild-type (wt) and *3h1* mutant seedlings grown for 19 days under long-day (16h light – 8h night) conditions, with the temperature fluctuating between 20^°^C at night and 22^°^C during the day. We profiled triplicate samples at six time points: ZT4, ZT13, ZT16, ZT20, ZT22, ZT24. These time points correlated with epigenomic changes at morning- or evening-phase loci (H3K9Ac at ZT4 and ZT16; H3K4me3 at ZT13) (data from Song et al, 2019) and peak expression of chromatin modifiers such as JMJ14 at ZT20, and SDG2 at ZT22 (Song et al., 2019). The replicate transcriptome profiles showed a high correlation at each time point (**Supplementary Figure S1C**). We then focused on genes with diurnal fluctuations and differential expressions between *3h1* and wt plants (LFC>1, FDR<0.01, n=1040, Time and Treatment Contrast, TTC genes, **Supplementary Data1**). Among them, we identified 34 genes specifically upregulated during lateral root development (Gala et al., 2021) (**Supplementary Data1**) which could explain the difference in lateral root number under continuous light. To gain a broader understanding of possible alteration in the diurnal kinetic of gene expression in *3h1*, we clustered the previously selected TTC genes according to their kinetics in the wt and compared their diurnal expression profile in the mutant (**Figure 1E**, 6 clusters; **Supplementary Figure S1D**, 8 clusters). In the wild type, gene clusters showed synchronised expression, whereas in the mutant, their expression was more variable and desynchronized for a subset of genes, peaking at different times compared to the wild type. In wild-type plants, H1-encoding genes do not exhibit diurnal changes in expression levels (**Supplementary Figure S1E**) suggesting that the impact of H1 depletion on diurnal gene regulation is not directly related to global H1 protein abundance. To further analyse genes most dramatically affected in their diurnal kinetics in the mutant, we used a deviation score that measures the cumulative discrepancy between wt and mutant profiles (**Supplementary Figure S1F**). Using this score, we selected genes with deviation scores in the top 5% range and bottom 5% range, respectively. This generated two gene groups, corresponding to de-synchronized genes (DSG, n=197) and synchronized genes (SG, n=773), respectively (**Figure 1F**).

### H1 depletion mostly affects PRC2 targets enriched in NAC binding motifs

To determine whether DSG genes, which show altered diurnal kinetics upon H1 depletion, are clock-regulated, we verified their expression levels in conditions where seedlings were first entrained by a photo- and a thermoperiod before being transferred to continuous light (DIURNAL data resource (Mockler et al., 2007)). The analysis showed that none of the DSG is regulated by the clock albeit two genes encoding subunits of the light harvesting complex (**Supplementary Figure S2A**). Instead, about one third of DSG are light-responsive according to a recent analysis of Arabidopsis seedling development s (Schivre et al., 2025) (**Supplementary Figure S2A**).

Next, to answer the question whether DSG represent a random or specific subset of diurnally regulated processes, we analysed the GO terms associated with DSG and SG. Nearly 80% of the biological processes enriched in DSG were linked to responses to biotic and abiotic stresses, compared to only 10% in SG (**Figure 2A**). In contrast, SG were enriched in fundamental cellular metabolism and nuclear functions (**Figure 2A**). This suggests that H1 regulates the timing of stress-response gene expression. DSG were generally expressed at lower levels than stably expressed genes (SGs) across all time points (**Figure 2B, Supplementary Figure S2B**). This lower expression correlated with higher average levels of H3K27me3 at DSG, at least at day start, based on available data (Baerenfeller et al., 2016; **Figure 2C, Supplementary Figure S2C**). While DSG showed similar average H1 enrichment to SGs (based on seedling data; time point not specified, Bourguet et al., 2021, **Figure 2C, Supplementary Figure S2C**), H3K27me3 levels at DSG may still be influenced by H1 occupancy. Supporting this, approximately half of the DSG overlap with regions depleted of H3K27me3 in cotyledon tissues of the *h1* double mutant (*h1*.*1 h1*.*2*) (Teano et al., 2023; **Supplementary Figure S2C**). Conversely, DSG show a significant depletion of H3K4me3, not only at day start (**Figure 2C**) but also across the diurnal phase (**Supplementary Figure S2D**, data from Song et al., 2019). Since H1 deposition interplays with DNA methylation (Wierzbicki & Jerzmanowski, 2005; Zemach et al., 2013), we also asked whether DSG were distinct in their DNA methylation levels compared to SG. Using available whole genome bisulfite sequencing data from 2 weeks old seedlings grown under a long day photoperiod (Rutowicz et al., 2015) we found that DSG were, in average, 8x less methylated in CpG context than SG (**Figure 2C**). In the CHG and CHH contexts, DSG exhibit similar methylation levels than SG in average albeit for a few loci showing elevated mCHG levels (**Supplementary Figure S2D**). Yet, DNA methylation levels do not change significantly more in DSG compared to SG in a *3h1* mutant background (**Supplementary Figure S2D**). This suggests that DNA methylation may play little role in the deregulation of DSG diurnal expression in *3h1* mutant seedlings.

**Figure 2.**
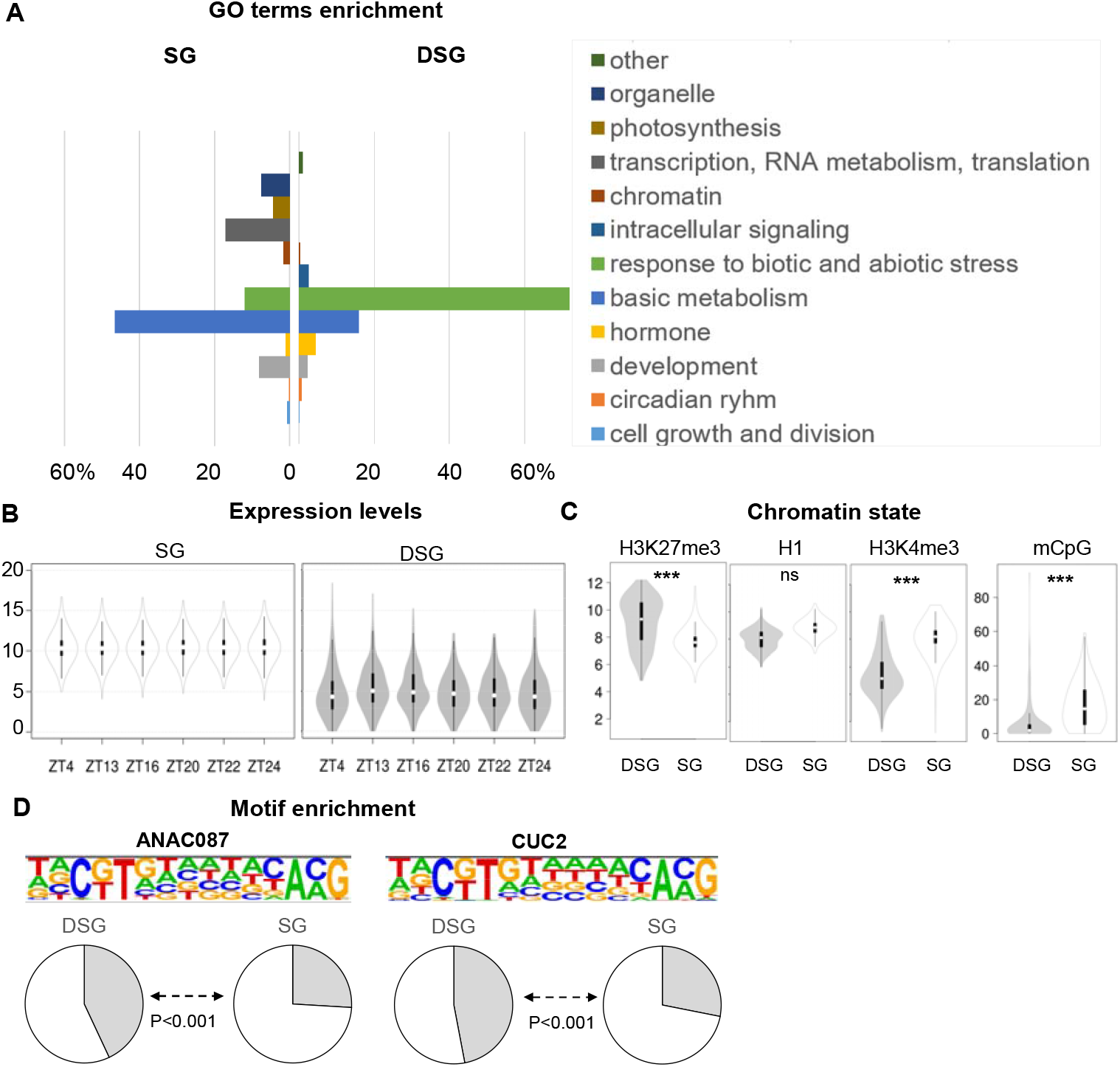
DSG are mostly response genes likely regulated by NAC transcription factors. **(A)** Genome Ontology analysis of DSG and SG showing biological processes that are significantly enriched (P<0.05). Processes are grouped in major categories indicated in the legend. The X axis shows the % of genes falling into each category. **(B)** Averaged, normalised expression levels of SG and DSG in the wild-type at each time point. **(C)** Relative enrichment levels of H3K27me3 at day start (Baerenfeller et al., 2016), H3K4me3 at ZT3 (Song et al. 2019), of H1 (Bourguet et al., 2021) and mCpG (Rutowicz et al., 2015) at unknown time points, in DSG vs SG. P values are from a Kruskal-Wallis test with Bonferroni correction. **(D)** Top two most enriched motifs among DSG and their occurrence in % among DSG vs SG (pie charts). See also **Supplementary Figure S2** and **Supplementary Data 2**.

Next, to investigate whether DSG regulation shares a common control mechanism, we searched for binding motifs of transcriptional regulators. We found a significant recurrence of motifs corresponding to NAC protein binding sites (**Figure 2D, Supplementary Data 2**). NAC proteins (NO APICAL MERISTEM, ATAF1/2, and CUP-SHAPED COTYLEDON) are part of a large, ancient, plant-specific transcription factor family involved in plant growth, senescence, flowering, fruit ripening, immunity, and abiotic stress responses (Kim et al., 2024) (Olsen et al., 2005). A role for NAC transcription factors in diurnal regulations has not been reported so far. While some of them can be diurnally or clock-regulated (Ma et al., 2025; Peng et al., 2020), it is not the case for our candidates whose binding motifs are enriched in DSGs (**Supplementary Figure S2E**). Whether NAC TFs interact with or compete against H1 proteins at binding sites to influence DSG rhythmic transcription remains to determine. Remarkably, DSG did not contain binding sites for circadian clock regulators, further supporting the idea that their rhythmic expression may be independently regulated or only indirectly controlled by the clock.

### H1 contributes to diurnal chromatin dynamics

H1 is broadly distributed across the genome (Bourguet et al., 2021; Rutowicz et al., 2015), and acts as a general chromatin organizer (Choi et al., 2020; He et al., 2024; Rutowicz et al., 2019; Teano et al., 2023). Despite this broad role, it can influence the transcriptional timing of specific genes (results above). We therefore asked whether H1 contribute large-scale chromatin reorganization that could possibly influence or accompany transcriptional reprogramming.

H1 visibly affects chromatin organization at the cytological level (Choi et al., 2020; He et al., 2024; Rutowicz et al., 2019) and 3D genome topology (Teano et al., 2023). Using quantitative imaging, we measured several nuclear and chromatin features. Nuclei were isolated from wild-type and *3h1* mutant seedlings at the same time points as before in replicate experiments (see Methods and **Supplementary Data 3**). In total, ~1530 DNA-stained nuclei were imaged (**Supplementary Figure S3A**). Using Nucl.Eye.D (Johann To Berens et al., 2022) we performed automated segmentation to quantify nuclear size (area), shape (roundness, circularity, solidity, aspect ratio), and DNA intensity distribution. While analysing the mean values of these features indicated variation across time points (**Supplementary Figure S3B, C**). A principal component analysis (PCA) best captured the diurnal changes, showing distinct trajectories during the day and night in wt nuclei (**Figure 3A**). Clearly, *3h1* mutant nuclei occupied a separate region in the PC space, likely due to their larger size and reduced heterochromatin content (Choi et al., 2020; Rutowicz et al., 2019), which also affects DNA intensity distribution (**Supplementary Figure S3A**). Yet, diurnal cytological changes showed a different trajectory in the mutant indicating that H1 plays a role in chromatin reorganization dynamics during the photoperiod (**Figure 3A**). We measured specifically the relative heterochromatin fraction (RHF) present in conspicuous chromocenters (Fransz et al., 2002). We observed that the RHF remained constant in average across time points and as expected (Rutowicz et al., 2019), the *3h1* mutant showed lower RHF and fewer chromocenters (**Figure 3B** and **Supplementary Figure S3D**).

**Figure 3.**
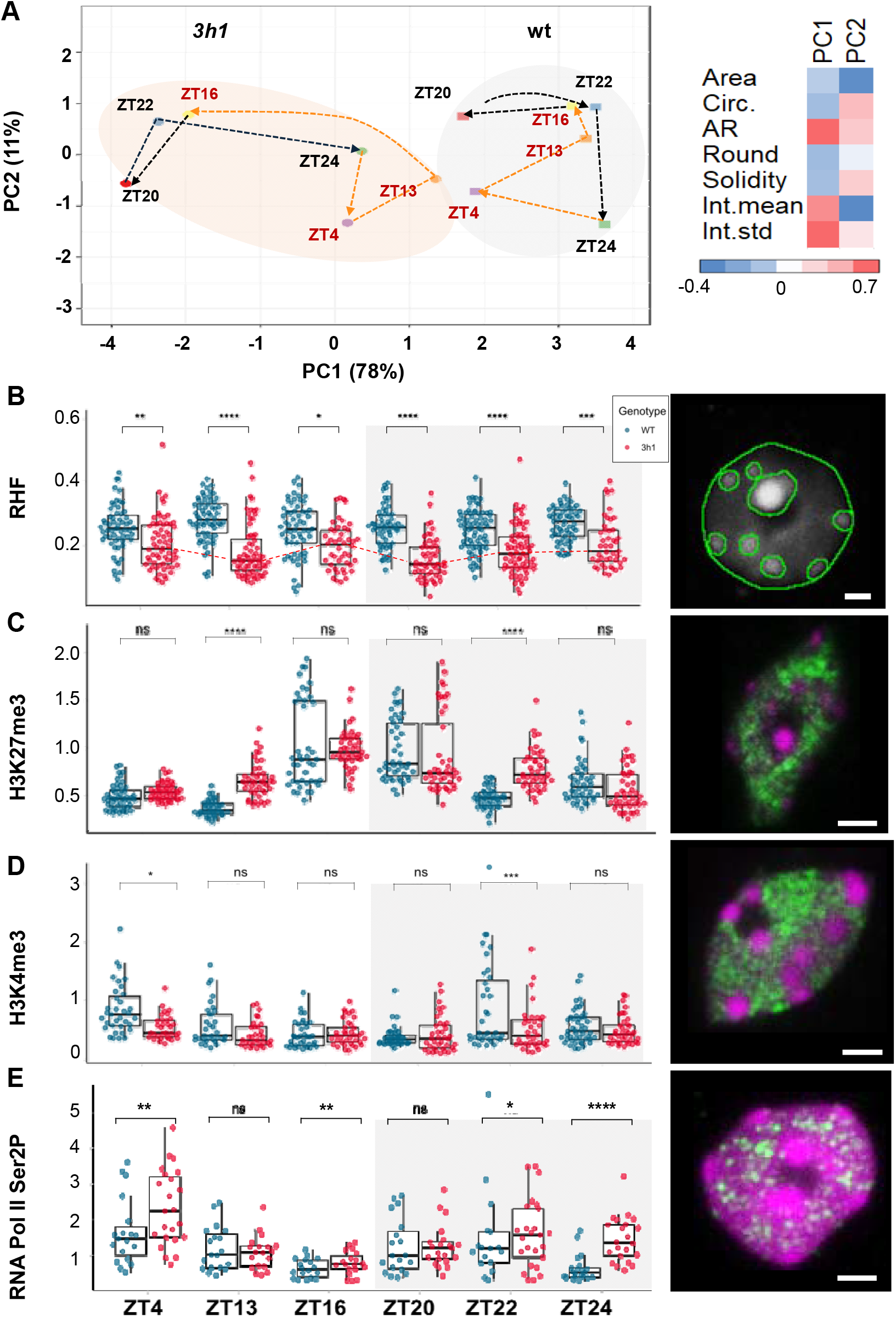
Diurnal changes in nuclear size, chromatin density, global epigenetic and transcriptional levels in Arabidopsis seedlings. Seedling nuclei were analysed by quantitative imaging following isolation, immunostaining and DNA counterstaining, before batch-segmentation using Nucl.Eye.D (Johann To Berens et al., 2022). (**A**) PCA analysis of nuclear features including area, shape descriptors (roundness, circularity, solidity, aspect ratio) and DNA density distribution (intensity mean and standard deviation -std). The distinct PC space occupied by wt and *3h1* samples, as well as distinct trajectories along the day-night cycle were manually colored and indicated. PC plot: columns with similar annotations were collapsed by taking the median inside each group. Unit variance scaling was applied to rows and SVD with imputation was used to calculate principal components (PC) 1 and 2 (X and Y axis) that explain 78% and 11% of the total variance, respectively. The weight of each descriptor for PC1 and PC2 are shown on the right. The number of nuclei analysed are given in **Supplementary Figure S3A** and range between 40 and 73 per sample. (**B-E**) Quantification of the relative heterochromatin content (RHF, B) and relative levels of H3K27me3 (C), H3K4me3 (D) and RNA Pol II Ser2P (E) in wt (blue) and 3h1 (red) nuclei. Relative levels: for each nucleus, the immunostaining signal intensity was normalised by the DNA signal intensity. Differences between genotypes were evaluated using a Mann Whitney U test. ns, not significant. *, P<0.05. **, P<0.01. ***, P<0.001. ****, P<0.0001. The right panels show representative images for each experiment: a DNA-stained nucleus segmented by nucl.Eye.D (B, green contours), nuclei immunostained (green) and counterstained with propidium iodide (magenta). Scale bar: 2µm. See also **Supplementary Figure S3, Supplementary Data 3** and **BioStudies accession S-BIAD1915**.

Despite its strong influence on the epigenetic landscape, H1 depletion has a modest effect on transcription in Arabidopsis (Choi et al., 2020; Hsieh et al., 2016; Lyons & Zilberman, 2017; Rutowicz et al., 2019; Rutowicz et al., 2015; Teano et al., 2023; Wierzbicki & Jerzmanowski, 2005; Zemach et al., 2013). Since the photoperiod is associated with rhythmic shifts in histone marks such as H3K4me3, H3K9Ac, H3K27Ac, H3K27me3, and H3K27me1 at diurnally regulated loci (Baerenfaller et al., 2016; Fung-Uceda et al., 2024; Song et al., 2019) we asked whether H1 might contribute to some of these changes. To test this, we used quantitative immunostaining for measuring the nuclear abundance of H3K4me3 and H3K27me3—marking transcriptionally permissive and repressive states, respectively—as well as productive RNA Polymerase II (Ser2-phosphorylated, Ser2P). In nuclei of wild-type seedlings, all three markers showed diurnal variations (**Supplementary Figure S3E-G**). Notably, H3K27me3 levels peaked at the end of the day (ZT16) and decreased before the end of the night (ZT24). Interestingly, a subset of nuclei at ZT16 and ZT20 showed a nearly 4-fold increase compared to ZT4, suggesting that the amplitude of this response might be cell-type specific. The *3h1* mutant also showed diurnal fluctuations in H3K27me3 but with less sharp transitions than in the wt, leading to significantly elevated levels towards the end of the day (ZT13) and end of the night (ZT22), (**Figure 3C**).

Conversely, both H3K4me3 and RNA Pol II Ser2P reached their lowest levels at ZT16-ZT20 rising slowly during the night until ZT4 in wt plants (**Supplementary Figure S3F-G**). The reduction in productive RNA Pol II (Ser 2P) at the end of the day correlated with a decrease in paused RNA Pol II (Ser 5P) (**Supplementary Figure S3H**), suggesting a global reduction in transcriptional activity as the night is approaches. In comparison, *3h1* nuclei also showed diurnal fluctuations but – as for H3K27me3 - with a strong attenuation of the diurnal variations. This led to higher levels of productive RNA Pol II (Ser2P) towards the end of the night (ZT22, ZT24) and lasting until the morning (ZT4) in conjunction with slightly elevated levels of H3K4me3 (**Figure 3D-E**).

To determine whether the increased RNA Pol II activity in *3h1* was a result from a reduction in the inactive fraction of RNA Pol II or reflected a general increase in available RNA Pol II, we quantified RNA Pol II Ser5P and total RNA Pol II (NP) levels, respectively, by immunostaining as before and from ZT4 to ZT16. The results supported the latter scenario (**Supplemental Figure I**).

In conclusion, our quantitative immunostaining analyses revealed large-scale, diurnal changes in chromatin organization and epigenetic composition, with fluctuations in global levels of H3K4me3 and H3K27me3 in wild-type seedlings. These changes are coincidental with a global reduction in RNA Pol II toward the end of the day, followed by an increase in the morning. In this process, linker histones appear to play a role, particularly in suppressing transcriptional activity during the night and early morning.

## Discussion

H1 linker histones are essential components of chromatin. They regulate its structure, compaction, and epigenetic landscape in both plants and animals (Fyodorov et al., 2018; He et al., 2024; Over & Michaels, 2014; Rutowicz et al., 2019; Rutowicz et al., 2015; Zemach et al., 2013). In Arabidopsis, H1 depletion causes mild developmental changes (Rutowicz et al., 2019). Our study revealed that H1 plays a role in synchronizing the expression of a subset of rhythmically regulated genes, particularly those involved in stress responses. This aligns with several studies highlighting the involvement of H1 variants in responses to biotic and abiotic stresses. For example, compared to wild-type plants, H1-depleted *Arabidopsis* exhibit enhanced resistance to *Pseudomonas syringae* and *Botrytis cinerea*, as well as increased defence priming in response to flg22 (Sheikh et al., 2023). They also show greater resistance to heat stress, particularly when combined with DNA methylation inhibition (Liu et al., 2021), but display hypersensitivity to salt stress (Perrella et al., 2024). Additionally, plants lacking the H1.3 variant demonstrate increased tolerance to drought and low-light conditions (Rutowicz et al., 2015)

While it is already known that a significant fraction of diurnally regulated genes are associated with shifts in various histone marks (Baerenfaller et al., 2016; Fung-Uceda et al., 2024; Song et al., 2019), our findings extend this knowledge by identifying rhythmic changes beyond these loci. Specifically, we observed changes in chromatin organization and nuclear levels of the key epigenetic marks H3K27me3 and H3K4me3, as well as in RNA Pol II levels. Additionally, we found that H1 contributes to these global changes. Given that H1 interacts closely with DNA methylation (Choi et al., 2020; Rutowicz et al., 2015; Wierzbicki & Jerzmanowski, 2005; Zemach et al., 2013), it remains to be determined whether global DNA methylation levels also fluctuate in a diurnal manner.

Several mechanisms, potentially cooperating, could explain the disrupted transcription timing in the absence of H1: a direct effect on RNA Pol II progression, an indirect effect through modulation of H3K4me3, H3K27me3 or other marks accumulation, such as DNA methylation, or a combination thereof. Indeed, H1 regulates RNA Polymerase II (RNA Pol II) progression during key phases of transcription. In both yeast and mammals, linker histones cause RNA Pol II to pause near the nucleosome entry site (Hirano et al., 2022). This is due to the close spatial proximity of the entry and exit DNA strands constrained by H1 on the nucleosome that creates a steric clash between the DNA exit strand and RNA Pol II, which is engaged with the entry strand. This conflict is resolved by transcription elongation factors, which remove H1 from the entry strand (Hirano et al., 2022). In Arabidopsis, the absence of H1 leads to irregular nucleosome spacing, with increased occurrences of both short (<170bp) and long (>300nt) nucleosome repeat lengths (NRL) (Choi et al., 2020; Rutowicz et al., 2019). Short nucleosome spacing may intensify steric clashes with RNA Pol II, that elongation factors cannot efficiently resolve. Conversely, longer NRL regions may promote extended RNA Pol II progression, where H1 would normally limit it. Thus, H1 absence could disrupt the regulation of transcriptional elongation during diurnal fluctuations. Also, intergenic transcripts, which emerge in the absence of H1 (Choi et al., 2020) is not excluded in contributing transcriptional timing. Additionally, in Arabidopsis, H1 is known to influence the deposition and maintenance of DNA methylation as well as histone marks such as H3K4me3 and H3K27me3 (Rutowicz et al., 2019; Rutowicz et al., 2015; Wierzbicki & Jerzmanowski, 2005; Zemach et al., 2013). Hence, the disrupted transcriptional timing observed in H1-deficient plants may be linked to altered epigenetic states that are poorly regulated in the absence of H1. Supporting this hypothesis, De-Synchronised Genes (DSG) in the *3h1* mutant are largely enriched in H3K27me3 and half display reduced H3K27me3 level upon H1 depletion while their expression increases in the absence of H1. DNA methylation on the contrary may not contribute much to DSG regulation since mCpG levels are much lower at DSG loci than SG and barely change upon H1 depletion. However, this alone does not fully explain changes in transcription kinetics. Temporally resolved, diurnal profiles of H3K27me3, and DNA methylation in both wild-type and *3h1* mutant plants are needed. Likely, however, additional factors may act in concert with H1 to integrate epigenetic states and enable diurnal transcriptional dynamics.

Linker histones (H1) are not essential for plant growth or viability under controlled conditions. However, under stress—both biotic and abiotic—H1 plays a role in modulating the stress response (Liu et al., 2021; Perrella et al., 2024; Rutowicz et al., 2015; Sheikh et al., 2023), influencing physiological and growth adaptations possibly through their influence on RNA Pol 2 and/or PRC2 activity (Teano et al., 2023). It is interesting to note that genes which diurnal regulation is disrupted upon H1 depletion are primarily involved in biotic and abiotic stress responses, heavily marked by H3K27me3 during 24-h cycles, and not regulated by the circadian clock. Clock-controlled genes, in contrast, are not subject to H3K27me3 deposition (Malapeira et al., 2012) and, often relate to core metabolic processes, light perception, and growth regulation although an interplay with abiotic responses are identified (Panter et al., 2019). The specificity of functional categories impacted by the absence of H1 is intriguing, especially considering the widespread deposition of H1 variants across the genome (Bourguet et al., 2021; Rutowicz et al., 2015). One plausible mechanism enabling selective effects of H1 at DSG is a competitive interaction between H1 and other transcriptional regulators, such as NAC transcription factors, which control stress responses and other processes (Olsen et al., 2005). This model is supported by the observation that NAC binding motifs are enriched at loci with desynchronized expression in the *3h1* mutant. In this context, H1 may restrict NAC factor access to these motifs. However, NAC target genes are not simply overexpressed throughout the photoperiod in the absence of H1 and their corresponding, candidate NAC transcription factors are themselves not diurnally regulated. This suggests that other factors modulate the competitive binding between H1 and NAC in a diurnal manner. One possibility is the role of H1 post-translational modifications (PTMs), which influence H1 binding kinetics (Catez et al., 2006). Phosphorylation is a prime candidate for destabilizing H1 from chromatin, which is especially relevant considering the significant fluctuations in the Arabidopsis phospho-proteome during the photoperiod (Uhrig et al., 2021). Competitive binding between H1 and specific chromatin regulators has been demonstrated, which explains the impact of H1 depletion at TRB target loci (Teano et al., 2023). Thus, this model of competitive binding suggests that H1, a generic chromatin factor, could regulate specific subsets of loci at certain times, potentially in response to environmental or developmental signals.

In conclusion, H1 may facilitate transcriptional reprogramming by coordinating chromatin accessibility, RNA Pol II progression, and competition with transcriptional regulators. However, the mechanisms by which these processes achieve precise temporal control at specific loci remain unclear and further research is required to elucidate these questions. Photoreceptors, metabolic feedback loops, or both may directly or indirectly influence the interaction between H1, epigenetic marks, NAC transcription factors, and RNA Pol II, ultimately synchronizing the expression of gene subsets with similar transcriptional kinetics.

## Material and Methods

### Plant growth

Arabidopsis thaliana Col_0 seedlings and Arabidopsis *3h1*^*crispr*^ mutant seedlings (**Supplemental File 1**) were grown on 0.5x Murashige & Skoog (MS) salt base (Carolina Biological Supply, US) with 1% PhytoAgar (Duchefa Biochemie, Netherlands) under continuous light or long-day conditions (16h/8h) depending on the experiment. The temperature was maintained at 21-22^°^C during the day and 18^°^C at night. For transcriptomic experiments, all seedlings were grown simultaneously in the same growth chamber, with six replicate plates per genotype and time point, randomly distributed on the shelves for sample collection. For cytological experiments, seedlings were grown in two separate incubators with an inverted day-night cycle to allow tissue collection for all time points within a day. Samples were collected in two replicate experiments (A and B) per genotype and time point.

### Quantification of root length and lateral root number

Seedlings were grown as before but vertically on square petri dishes and scanned 7 and 12 days after germination before measurements of root length using Fiji (Schindelin et al., 2012) and manual scoring of lateral roots.

### RNA-seq data processing

Short reads generated in this study were deposited at NCBI Sequence Read Archive (SRA, www.ncbi.nlm.nih.gov/sra) and are accessible through the accession number <u>PRJNA1244618</u>. Reads were trimmed and quality-checked with fastp (Chen et al., 2018) (version 0.20.1). Transcripts were quantified with Salmon (Patro et al., 2017) (version 1.4.0) using the cDNA and gene annotation available from araport.org (Araport 11). Variation in read counts was analyzed with a general linear model in R (version 3.6.1) with the package DESeq2 (Love et al., 2014) (version 1.24.0) according to a factorial design with the two explanatory factors “time” and “treatment” combined into a single factor. Specific conditions were then compared with linear contrasts. Within each comparison, *P*-values were adjusted for multiple testing (Benjamini-Hochberg). Regions with an adjusted *P*-value (false discovery rate, FDR) below 0.01 and a minimal log2 fold-change (i.e., the difference between the log2 transformed, normalized sequence counts) of 1 were differentially expressed. Normalized sequence counts were calculated accordingly with DESeq2 and log2(x+1) transformed.

### Definition of gene sets

To identify genes with differences in diurnal expression between the *3h1* mutant and the wild-type, we used 1) linear, quadratic, and cubic contrast (Rosenthal & Rosnow, 1985); 2) contrasts that compare one time point to all other time points (one test for each other time point); and 3) contrasts that compare two adjacent time points to all other time points (also one test for each other time point). Genes that showed significant differences in any of these contrasts using the wild-type data were considered “genes that change over time”.

To identify genes with differences in expression between *3h1* and wild-type, the treatments were compared at each and across all time points. Genes that showed significant differences in any of these comparisons were defined as “genes that change by treatment”. However, to focus on genes that show the most “de-synchronized” pattern, we calculated an index of change as the sum of the absolute LFCs between *3h1* and wt at all individual time points. Large values indicate that genes show a *3h1*-specific pattern that is clearly distinct from the wild-type and not just shifted in intensity. From all genes, the top 2.5 % was intersected with the genes that change by treatment, resulting in 216 de-synchronized genes. Likewise, we devised a set of genes that were synchronized between *3h1* and wt seedlings by intersecting the bottom 2.5 % with the genes that don’t show any differences in expression between treatments (578 synchronized genes).

### Time course visualization

To visualize the time-course behaviour of genes of interest, we first grouped genes into clusters using wild-type data. For this, replicates from each time and treatment combination were averaged. Expression of each gene was standardized and the wild-type data was clustered into 4 to 8 clusters using Mfuzz (Kumar & M, 2007). *3h1* data was added to the visualization but not used while clustering. Only genes belonging to the cluster core are shown. Those were extracted with the function acore() from the Mfuzz package (Kumar & M, 2007).

### Functional gene set characterization

To functionally characterize a gene set of interest (e.g., synchronized and de-synchronized genes), we tested for enrichment of gene ontology (GO) terms with topGO (version 2.32.0) (Alexa et al., 2006). Analysis was based on gene counts (genes in the set of interest compared to all annotated genes) using the “weight” algorithm with Fisher’s exact test (both implemented in topGO). A term was identified as significant if the *P*-value was below 0.05.

### Motif discovery

To search for motifs in promoters (2 kb upstream of TSS) of genes of interest, we used findMotifs.pl from the HOMER suite (version 4.11) (Heinz et al., 2010) with the synchronized genes as control group. As we observed many NAC matches, we extracted all matches and generated a consensus motif with the R-package “<u>universalmotif</u>” (https://github.com/bjmt/universalmotif). The consensus motif was then searched in the promoter sequences with fimo (meme suite, version 5.0.2, 2011) (Bailey et al., 2009).

### Cross-analyses with ChIP-seq and RNA-seq data

Publicly available ChIP-seq raw data were retrieved from SRA/NCBI. Datasets from seedling tissues included H3K4me3 and H3K9me3 (Song et al., 2019), H3K27me3 (Veluchamy et al., 2016) and H1 (Bourguet et al., 2021). Reads were trimmed and quality-checked with fastp (version 0.20.1) (Chen et al., 2018) and aligned to the Arabidopsis reference genome (TAIR10) with bowtie 2 (version 2.3.5.1) (Langmead & Salzberg, 2012) keeping only unique alignments. Duplicate reads were removed with Picard (version 1.140; <u>broadinstitute.github.io/picard/</u>). Coverage was normalized and visualized with DeepTools (version 3.3.0) (Ramirez et al., 2014); bamCoverage with --normalizeUsing CPM, computeMatrix scale-regions, and plotHeatmap). In addition, determination of differential H3K27me3 enrichment in wt and *2h1* seedlings for DSG and SG was done according to a DEseq2 analysis reported in [Teano et al 2023, Table S1). The values reported in Supplemental Table 2 represent Log2 fold change, with NA corresponding to genes without significant H3K27me3 enrichment upon peak detection with MACS2. The cross-analysis of DSG and SG with light-induced genes during cotyledon de-etiolation was done using a Spike-In RNAseq dataset (Schivre et al., 2025). Values represent normalised transcript levels, with NA corresponding to genes with no detectable reads.

### WGBS data processing

Short reads from whole genome bisulfite sequencing for Col_0 wt seedlings and *3h1* (T-DNA allele, Rutowicz et al. 2015) mutant seedlings were obtained from ArrayExpress (E-MTAB-2807). Reads were (quality) trimmed with fastp (version 0.20.0 with the options --trim_front1 2, (Chen et al., 2018) and aligned to the reference genome with Bowtie2 (version 2.3.5.1, (Langmead & Salzberg, 2012) in combination with Bismark (version 0.22.3 with the option --non_directional (Krueger & Andrews, 2011)). The Col-0 reference genome was obtained from www.arabidopsis.org (TAIR10). Alignments were deduplicated with bismark and methylation tables were extracted with MethylDackel (version 0.5.0, github.com/dpryan79/MethylDackel). Files were merged and only cytosines with a coverage between 5 and 100 within all four samples were kept. 11.1 million cytosines passed this filter. Cytosines were mapped to the Araport 11 annotation to obtain average methylation levels per gene and methylation context. For each gene and context, we averaged the methylation levels for the two groups (wt and *3h1*) and calculated the difference between the groups (delta). Genes that were identified as SG and DSG were extracted and were tested within each context for different DNA methylation levels in wt comparing SG and DSG groups, or differential response in *3h1* vs wt (delta) using a two-sided t-test.

To assess differences in DNA-methylation in more detail, the two groups (3h1 and wt) were compared to each other with the R-package DSS (version 2.32.0, (Park & Wu, 2016)) using the functions DMLfit.multiFactor() and DMLtest.multiFactor(). P-values were corrected for multiple testing and thereby converted to false discovery rates (FDRs, (Benjamini & Hochberg, 1995). Cytosines were identified as significantly differentially methylated (DMC) if the FDR was below 0.05. 25,218 cytosines exhibited significant differential methylation (0.23 %). We extracted the genes for which there were at least 10 cytosines in one of the contexts tested (34,626) and subset them to the ones with at least 20 % differentially methylated cytosines: 468 genes were left but only 10 of them were in the list of SGs and DSGs. However, all of them were SGs. The genes were AT1G31440, AT2G03350, AT2G04540, AT2G04880, AT2G14120, AT2G14680, AT2G17410, AT2G18876, AT2G19430, and AT2G19470.

### Nuclei extraction and immunostaining

Nuclei were extracted essentially as described previously (Kracik-Dyer & Baroux, 2025), leaving out the embedding step and adding a clearing procedure. In brief, leaves from 19-day-old *A. thaliana* seedlings were fixed, washed then finely chopped with a sterile razor blade in nuclear isolation buffer. The suspension was briefly sonicated and filtered before distributed as 5_µ_L drops on cleaned slides (Epredia^Tm^ SuperFrost Plus^TM^ Adhesion Slides, ThermoFisher Scientific). Nuclei were air-dried until used or stored at 4^°^C. Prior to immunostaining, slides underwent a clearing treatment to enhance downstream imaging quality. Specifically, slides were placed in a Coplin jar and treated for 5 minutes in a 1:1 solution of 100% ethanol and xylene, followed by a 5-minute wash in 100% methanol, a 5-minute wash in a 1:1 solution of methanol and 1× PBS, and a final 10-minute wash in 1× PBS.

Immunostaining was carried out following a 45-minute blocking step at room temperature using 4% BSA in 1xPBS,143 mM NaCl, followed by a 10-minute wash in 1× PBS. The primary and secondary antibodies were each applied sequentially at a 1:1000 dilution after having verified that this allowed a quantitative analysis (**Supplementary Figure S3J**) in 1× PBS containing 1% BSA, 143 mM NaCl, 0.05% Tween20 and incubated at 37□ ^°^C for 30 minutes each, with a 10-minute wash in 1× PBS between applications. Antibodies used in this study: Rabbit-anti-H3K27me3 (Active Motif cat# 39155), Rabbit-anti-H3K4me3 (Abcam cat# ab8580), Rabbit-anti-RNA Polymerase II CTD Phospho S2 (Abcam cat# ab5095), Rabbit-anti-RNA Polymerase II CTD Phospho S5 (Abcam cat# ab5131), Goat-anti-Rabbit Alexa 488 conjugate (ThermoFisher Scientific, cat# A-11008). Nuclei were stained with 2µg/mL 4_′_,6-diamidino-2-phenylindole (DAPI) or 2µg/mL Propidium Iodide (PI) in PBS for 10min, washed 1 min in PBS and mounted in VECTASHIELD® Antifade Mounting Medium complemented with DAPI (Vector Laboratories H-1100) or PI (Vector Laboratories H-1300), respectively, before being covered with 18×18mm, 0.17mm thick cover glasses, sealed with transparent nail polish and stored in the dark at 4^°^C until imaging.

### Fluorescence Microscopy Imaging

To minimize selection bias, nuclei were consistently selected based on DNA staining with DAPI rather than antibody staining. Nuclei that appeared damaged or morphologically abnormal were excluded from imaging. Images were acquired using a Leica DM6000B microscope (Leica microsystems GmbH, Germany) equipped with a 63×/1.4 NA oil immersion objective and an Andor Neo 5.5 sCMOS camera (Andor, Oxford Scientific Instruments, USA). Images were captured in a 16-bit format using the ‘low-noise gain’ setting, and 15-200ms exposure, depending on the antibody and fluorescence filters (Chroma Technology Corp, USA: A4, L5 and Tx2 for DAPI, Alexa488 and PI detection, respectively). For quantifying immunostaining signal, single-plane images were acquired, while short z-stacks were acquired -using system’s optimized settings-for downstream quantification of the relative heterochromatin fraction (RHF). Acquisition parameters were set for each fluorochrome to achieve a good pixel intensity distribution while avoiding saturation and kept constant throughout the slides.

### Image analysis

For RHF analyses, images were segmented and quantified using Nucl.eye.D using the author’s instructions (Johann To Berens et al., 2022) and using 2D projections of the z-stacks as input images. Max projections were created in batch using a macro in Fiji (Schindelin et al., 2012). The quantification of immunostaining signals was carried out using Fiji (Schindelin et al., 2012). For this, a macro was written to (i) separate the channels for each image (Image/Color/Split Channels), (ii) stack all images in one file (Image/Stacks/Images to Stack/Copy (center); Use Titles as Labels), create one file per channel, (iii) create a montage for each stack file (Image/Stacks/Make Montage using Increment=2, 6 Columns, 3 Rows – adjust to needs, First slice=1 for the first montage, first channel; First slice=2 for the second montage, second channel). Then, Regions of Interest (ROIs) were manually drawn around each nucleus in the montage containing the DNA signal, with ROIs saved in the ROI manager and used to extract signal intensities. The saved ROIs were opened and applied to the other montage to extract signal intensities from the second channel. Signal intensities were expressed as a ratio (Antibody to DNA) for the creating the graphs, plotted with RStudio (RStudio [2022.12.0+353] http://www.rstudio.com/). Unless otherwise stated, the data distribution across timepoints and genotypes were statistically evaluated using an ANOVA with Tukey HSD test. PCA analyses were conducted with the online tool ClustVis (Metsalu & Vilo, 2015).

## Acknowledgements

The authors thank Valeria Gagliardini, Christoph Eichenberger, Arturo Bolanos and Peter Kopf (University of Zurich) for general lab support, Professor Ueli Grossniklaus (University of Zürich) for valuable discussions.

## Data and Coding Availability Statement

RNAseq data are available at NCBI Sequence Read Archive (SRA) with the reference number PRJNA1244618. Image data are available at Biostudies accession number S-BIAD1915.

## Author contributions

CBa, KR conceived and designed the study, PZ and AJ designed elements of the study. KR, LP, JS, RSR, DZP, TB, MK performed the experiments. MWM performed bioinformatic analyses. KR, LP, CBo, FB and CBa performed data analyses and statistical analyses. CB wrote the manuscript. KR, LP, PZ, TB, MK CBo, FB contributed manuscript amelioration.

## References

Alexa, A., Rahnenfuhrer, J., & Lengauer, T. (2006). Improved scoring of functional groups from gene expression data by decorrelating GO graph structure. Bioinformatics, 22(13), 1600–1607. doi:10.1093/bioinformatics/btl140

Baerenfaller, K., Shu, H., Hirsch-Hoffmann, M., Futterer, J., Opitz, L., Rehrauer, H., Hennig, L., & Gruissem, W. (2016). Diurnal changes in the histone H3 signature H3K9ac|H3K27ac|H3S28p are associated with diurnal gene expression in Arabidopsis. Plant Cell Environ, 39(11), 2557–2569. doi:10.1111/pce.12811

Bailey, T. L., Boden, M., Buske, F. A., Frith, M., Grant, C. E., Clementi, L., Ren, J., Li, W. W., & Noble, W. S. (2009). MEME SUITE: tools for motif discovery and searching. Nucleic Acids Res, 37(Web Server issue), W202–208. doi:10.1093/nar/gkp335

Balcerowicz, M. (2024). A night shift for histone methylation in DNA damage control. Plant J, 120(6), 2323–2324. doi:10.1111/tpj.17192

Benjamini, Y., & Hochberg, Y. (1995). Controlling the False Discovery Rate: A Practical and Powerful Approach to Multiple Testing. Journal of the Royal Statistical Society: Series B (Methodological), 57(1), 289–300. doi:10.1111/j.2517-6161.1995.tb02031.x

Bieluszewski, T., Szymanska-Lejman, M., Dziegielewski, W., Zhu, L., & Ziolkowski, P.A. (2022). Efficient Generation of CRISPR/Cas9-Based Mutants Supported by Fluorescent Seed Selection in Different Arabidopsis Accessions. Methods Mol Biol, 2484, 161–182. doi:10.1007/978-1-0716-2253-7_13

Bourguet, P., Picard, C. L., Yelagandula, R., Pelissier, T., Lorkovic, Z. J., Feng, S., Pouch-Pelissier, M. N., Schmucker, A., Jacobsen, S. E., Berger, F., & Mathieu, O. (2021). The histone variant H2A.W and linker histone H1 co-regulate heterochromatin accessibility and DNA methylation. Nat Commun, 12(1), 2683. doi:10.1038/s41467-021-22993-5

Catez, F., Ueda, T., & Bustin, M. (2006). Determinants of histone H1 mobility and chromatin binding in living cells. Nat Struct Mol Biol, 13(4), 305–310. doi:10.1038/nsmb1077

Chen, S., Zhou, Y., Chen, Y., & Gu, J. (2018). fastp: an ultra-fast all-in-one FASTQ preprocessor. Bioinformatics, 34(17), i884–i890. doi:10.1093/bioinformatics/bty560

Choi, J., Lyons, D. B., Kim, M. Y., Moore, J. D., & Zilberman, D. (2020). DNA Methylation and Histone H1 Jointly Repress Transposable Elements and Aberrant Intragenic Transcripts. Mol Cell, 77(2), 310–323 e317. doi:10.1016/j.molcel.2019.10.011

Coen, E., & Prusinkiewicz, P. (2024). Developmental timing in plants. Nat Commun, 15(1), 2674. doi:10.1038/s41467-024-46941-1

Deng, L., Gao, B., Zhao, L., Zhang, Y., Zhang, Q., Guo, M., Yang, Y., Wang, S., Xie, L., Lou, H., Ma, M., Zhang, W., Cao, Z., Zhang, Q., McClung, C. R., Li, G., & Li, X. (2022). Diurnal RNAPII-tethered chromatin interactions are associated with rhythmic gene expression in rice. Genome Biol, 23(1), 7. doi:10.1186/s13059-021-02594-7

Duncan, O., & Millar, A. H. (2022). Day and night isotope labelling reveal metabolic pathway specific regulation of protein synthesis rates in Arabidopsis. Plant J, 109(4), 745–763. doi:10.1111/tpj.15661

Fransz, P., De Jong, J. H., Lysak, M., Castiglione, M. R., & Schubert, I. (2002). Interphase chromosomes in Arabidopsis are organized as well defined chromocenters from which euchromatin loops emanate. Proc Natl Acad Sci U S A, 99(22), 14584–14589. doi:10.1073/pnas.212325299

Fung-Uceda, J., Gomez, M. S., Rodriguez-Casillas, L., Gonzalez-Gil, A., & Gutierrez, C. (2024). Diurnal control of H3K27me1 deposition shapes expression of a subset of cell cycle and DNA damage response genes. Plant J, 120(6), 2325– 2336. doi:10.1111/tpj.17114

Fyodorov, D. V., Zhou, B. R., Skoultchi, A. I., & Bai, Y. (2018). Emerging roles of linker histones in regulating chromatin structure and function. Nat Rev Mol Cell Biol, 19(3), 192–206. doi:10.1038/nrm.2017.94

Gala, H. P., Lanctot, A., Jean-Baptiste, K., Guiziou, S., Chu, J. C., Zemke, J. E., George, W., Queitsch, C., Cuperus, J. T., & Nemhauser, J. L. (2021). A single-cell view of the transcriptome during lateral root initiation in Arabidopsis thaliana. Plant Cell, 33(7), 2197–2220. doi:10.1093/plcell/koab101

He, S., Yu, Y., Wang, L., Zhang, J., Bai, Z., Li, G., Li, P., & Feng, X. (2024). Linker histone H1 drives heterochromatin condensation via phase separation in Arabidopsis. Plant Cell, 36(5), 1829–1843. doi:10.1093/plcell/koae034

Heinz, S., Benner, C., Spann, N., Bertolino, E., Lin, Y. C., Laslo, P., Cheng, J. X., Murre, C., Singh, H., & Glass, C. K. (2010). Simple combinations of lineage-determining transcription factors prime cis-regulatory elements required for macrophage and B cell identities. Mol Cell, 38(4), 576–589. doi:10.1016/j.molcel.2010.05.004

Hirano, R., Ehara, H., Kujirai, T., Uejima, T., Takizawa, Y., Sekine, S. I., & Kurumizaka, H. (2022). Structural basis of RNA polymerase II transcription on the chromatosome containing linker histone H1. Nat Commun, 13(1), 7287. doi:10.1038/s41467-022-35003-z

Hsieh, P. H., He, S., Buttress, T., Gao, H., Couchman, M., Fischer, R. L., Zilberman, D., & Feng, X. (2016). Arabidopsis male sexual lineage exhibits more robust maintenance of CG methylation than somatic tissues. Proc Natl Acad Sci U S A, 113(52), 15132–15137. doi:10.1073/pnas.1619074114

Johann To Berens, P., Schivre, G., Theune, M., Peter, J., Sall, S. O., Mutterer, J., Barneche, F., Bourbousse, C., & Molinier, J. (2022). Advanced Image Analysis Methods for Automated Segmentation of Subnuclear Chromatin Domains. Epigenomes, 6(4). doi:10.3390/epigenomes6040034

Jung, S., Kim, H., Lee, J., Kang, M. H., Kim, J., Kim, J. K., Lim, P. O., & Nam, H. G. (2024). The genetically programmed rhythmic alteration of diurnal gene expression in the aged Arabidopsis leaves. Front Plant Sci, 15, 1481682. doi:10.3389/fpls.2024.1481682

Kim, T., Alvarez, J. C., Rana, D., Preciado, J., Liu, T., & Begcy, K. (2024). Evolution of NAC transcription factors from early land plants to domesticated crops. Plant Cell Physiol. doi:10.1093/pcp/pcae133

Kracik-Dyer, E., & Baroux, C. (2025). 3D STED Imaging of Isolated Arabidopsis thaliana Nuclei. Methods Mol Biol, 2873, 263–280. doi:10.1007/978-1-0716-4228-3_15

Krueger, F., & Andrews, S. R. (2011). Bismark: a flexible aligner and methylation caller for Bisulfite-Seq applications. Bioinformatics, 27(11), 1571–1572. doi:10.1093/bioinformatics/btr167

Kumar, L., & M, E. F. (2007). Mfuzz: a software package for soft clustering of microarray data. Bioinformation, 2(1), 5–7. doi:10.6026/97320630002005

Langmead, B., & Salzberg, S. L. (2012). Fast gapped-read alignment with Bowtie 2. Nat Methods, 9(4), 357–359. doi:10.1038/nmeth.1923

Le Martelot, G., Canella, D., Symul, L., Migliavacca, E., Gilardi, F., Liechti, R., Martin, O., Harshman, K., Delorenzi, M., Desvergne, B., Herr, W., Deplancke, B., Schibler, U., Rougemont, J., Guex, N., Hernandez, N., Naef, F., & Cycli, X. C. (2012). Genome-wide RNA polymerase II profiles and RNA accumulation reveal kinetics of transcription and associated epigenetic changes during diurnal cycles. PLoS Biol, 10(11), e1001442. doi:10.1371/journal.pbio.1001442

Li, L., Duncan, O., Ganguly, D. R., Lee, C. P., Crisp, P. A., Wijerathna-Yapa, A., Salih, K., Trosch, J., Pogson, B. J., & Millar, A. H. (2022). Enzymes degraded under high light maintain proteostasis by transcriptional regulation in Arabidopsis. Proc Natl Acad Sci U S A, 119(20), e2121362119. doi:10.1073/pnas.2121362119

Liu, S., de Jonge, J., Trejo-Arellano, M. S., Santos-Gonzalez, J., Kohler, C., & Hennig, L. (2021). Role of H1 and DNA methylation in selective regulation of transposable elements during heat stress. New Phytol, 229(4), 2238–2250. doi:10.1111/nph.17018

Love, M. I., Huber, W., & Anders, S. (2014). Moderated estimation of fold change and dispersion for RNA-seq data with DESeq2. Genome Biol, 15(12), 550. doi:10.1186/s13059-014-0550-8

Lyons, D. B., & Zilberman, D. (2017). DDM1 and Lsh remodelers allow methylation of DNA wrapped in nucleosomes. Elife, 6. doi:10.7554/eLife.30674

Ma, C., Zhou, P., Ma, Y., Tan, W., Huang, X., Segbo, S., Iqbal, S., Shi, T., Ni, Z., & Gao, Z. (2025). A NAC family gene PmNAC32 associated with photoperiod promotes flower induction in Prunus mume. Hortic Res, 12(9), uhaf157. doi:10.1093/hr/uhaf157

Malapeira, J., Khaitova, L. C., & Mas, P. (2012). Ordered changes in histone modifications at the core of the Arabidopsis circadian clock. Proc Natl Acad Sci U S A, 109(52), 21540–21545. doi:10.1073/pnas.1217022110

Maric, A., & Mas, P. (2020). Chromatin Dynamics and Transcriptional Control of Circadian Rhythms in Arabidopsis. Genes (Basel), 11(10). doi:10.3390/genes11101170

Matthews, J. S. A., Vialet-Chabrand, S., & Lawson, T. (2018). Acclimation to Fluctuating Light Impacts the Rapidity of Response and Diurnal Rhythm of Stomatal Conductance. Plant Physiol, 176(3), 1939–1951. doi:10.1104/pp.17.01809

Metsalu, T., & Vilo, J. (2015). ClustVis: a web tool for visualizing clustering of multivariate data using Principal Component Analysis and heatmap. Nucleic Acids Res, 43(W1), W566-570. doi:10.1093/nar/gkv468

Mockler, T. C., Michael, T. P., Priest, H. D., Shen, R., Sullivan, C. M., Givan, S. A., McEntee, C., Kay, S. A., & Chory, J. (2007). The DIURNAL project: DIURNAL and circadian expression profiling, model-based pattern matching, and promoter analysis. Cold Spring Harb Symp Quant Biol, 72, 353–363. doi:10.1101/sqb.2007.72.006

Nozue, K., & Maloof, J. N. (2006). Diurnal regulation of plant growth. Plant Cell Environ, 29(3), 396–408. doi:10.1111/j.1365-3040.2005.01489.x

Olsen, A. N., Ernst, H. A., Leggio, L. L., & Skriver, K. (2005). NAC transcription factors: structurally distinct, functionally diverse. Trends Plant Sci, 10(2), 79– 87. doi:10.1016/j.tplants.2004.12.010

Over, R. S., & Michaels, S. D. (2014). Open and closed: the roles of linker histones in plants and animals. Mol Plant, 7(3), 481–491. doi:10.1093/mp/sst164

Panter, P. E., Muranaka, T., Cuitun-Coronado, D., Graham, C. A., Yochikawa, A., Kudoh, H., & Dodd, A. N. (2019). Circadian Regulation of the Plant Transcriptome Under Natural Conditions. Front Genet, 10, 1239. doi:10.3389/fgene.2019.01239

Park, Y., & Wu, H. (2016). Differential methylation analysis for BS-seq data under general experimental design. Bioinformatics, 32(10), 1446–1453. doi:10.1093/bioinformatics/btw026

Patro, R., Duggal, G., Love, M. I., Irizarry, R. A., & Kingsford, C. (2017). Salmon provides fast and bias-aware quantification of transcript expression. Nat Methods, 14(4), 417–419. doi:10.1038/nmeth.4197

Peng, H., Phung, J., Zhai, Y., & Neff, M. M. (2020). Self-transcriptional repression of the Arabidopsis NAC transcription factor ATAF2 and its genetic interaction with phytochrome A in modulating seedling photomorphogenesis. Planta, 252(4), 48. doi:10.1007/s00425-020-03456-5

Perales, M., & Mas, P. (2007). A functional link between rhythmic changes in chromatin structure and the Arabidopsis biological clock. Plant Cell, 19(7), 2111–2123. doi:10.1105/tpc.107.050807

Perrella, G., Fasano, C., Donald, N. A., Daddiego, L., Fang, W., Martignago, D., Carr, C., Conti, L., Herzyk, P., & Amtmann, A. (2024). Histone Deacetylase Complex 1 and histone 1 epigenetically moderate stress responsiveness of Arabidopsis thaliana seedlings. New Phytol, 241(1), 166–179. doi:10.1111/nph.19165

Ramirez, F., Dundar, F., Diehl, S., Gruning, B. A., & Manke, T. (2014). deepTools: a flexible platform for exploring deep-sequencing data. Nucleic Acids Res, 42(Web Server issue), W187-191. doi:10.1093/nar/gku365

Redmond, E. J., Ronald, J., Davis, S. J., & Ezer, D. (2024). Stable and dynamic gene expression patterns over diurnal and developmental timescales in <em>Arabidopsis thaliana</em>. bioRxiv, 2024.2009.2018.613638. doi:10.1101/2024.09.18.613638

Rosenthal, R., & Rosnow, R. L. (1985). Contrast Analysis: Focused Comparisons in the Analysis of Variance: Cambridge University Press.

Rutowicz, K., Lirski, M., Mermaz, B., Teano, G., Schubert, J., Mestiri, I., Kroten, M. A., Fabrice, T. N., Fritz, S., Grob, S., Ringli, C., Cherkezyan, L., Barneche, F., Jerzmanowski, A., & Baroux, C. (2019). Linker histones are fine-scale chromatin architects modulating developmental decisions in Arabidopsis. Genome Biol, 20(1), 157. doi:10.1186/s13059-019-1767-3

Rutowicz, K., Puzio, M., Halibart-Puzio, J., Lirski, M., Kotlinski, M., Kroten, M. A., Knizewski, L., Lange, B., Muszewska, A., Sniegowska-Swierk, K., Koscielniak, J., Iwanicka-Nowicka, R., Buza, K., Janowiak, F., Zmuda, K., Joesaar, I., Laskowska-Kaszub, K., Fogtman, A., Kollist, H., Zielenkiewicz, P., Tiuryn, J., Siedlecki, P., Swiezewski, S., Ginalski, K., Koblowska, M., Archacki, R., Wilczynski, B., Rapacz, M., & Jerzmanowski, A. (2015). A Specialized Histone H1 Variant Is Required for Adaptive Responses to Complex Abiotic Stress and Related DNA Methylation in Arabidopsis. Plant Physiol, 169(3), 2080–2101. doi:10.1104/pp.15.00493

Schindelin, J., Arganda-Carreras, I., Frise, E., Kaynig, V., Longair, M., Pietzsch, T., Preibisch, S., Rueden, C., Saalfeld, S., Schmid, B., Tinevez, J. Y., White, D. J., Hartenstein, V., Eliceiri, K., Tomancak, P., & Cardona, A. (2012). Fiji: an open-source platform for biological-image analysis. Nat Methods, 9(7), 676– 682. doi:10.1038/nmeth.2019

Schivre, G., Wolff, L., Mirasole, F. M., Armanet, E., Davidson, M. L. H., Vidal, A., Cuménal, D., Dumont, M., Bourge, M., Baroux, C., Bourbousse, C., & Barneche, F. (2025). Genome-scale transcriptome augmentation during Arabidopsis thaliana photomorphogenesis. bioRxiv, 2025.2001.2030.635720. doi:10.1101/2025.01.30.635720

Seaton, D. D., Graf, A., Baerenfaller, K., Stitt, M., Millar, A. J., & Gruissem, W. (2018). Photoperiodic control of the Arabidopsis proteome reveals a translational coincidence mechanism. Mol Syst Biol, 14(3), e7962. doi:10.15252/msb.20177962

Sheikh, A. H., Nawaz, K., Tabassum, N., Almeida-Trapp, M., Mariappan, K. G., Alhoraibi, H., Rayapuram, N., Aranda, M., Groth, M., & Hirt, H. (2023). Linker histone H1 modulates defense priming and immunity in plants. Nucleic Acids Res, 51(9), 4252–4265. doi:10.1093/nar/gkad106

Song, Q., Huang, T. Y., Yu, H. H., Ando, A., Mas, P., Ha, M., & Chen, Z. J. (2019). Diurnal regulation of SDG2 and JMJ14 by circadian clock oscillators orchestrates histone modification rhythms in Arabidopsis. Genome Biol, 20(1), 170. doi:10.1186/s13059-019-1777-1

Teano, G., Concia, L., Wolff, L., Carron, L., Biocanin, I., Adamusova, K., Fojtova, M., Bourge, M., Kramdi, A., Colot, V., Grossniklaus, U., Bowler, C., Baroux, C., Carbone, A., Probst, A. V., Schrumpfova, P. P., Fajkus, J., Amiard, S., Grob, S., Bourbousse, C., & Barneche, F. (2023). Histone H1 protects telomeric repeats from H3K27me3 invasion in Arabidopsis. Cell Rep, 42(8), 112894. doi:10.1016/j.celrep.2023.112894

Tian, H., Li, Y., Wang, C., Xu, X., Zhang, Y., Zeb, Q., Zicola, J., Fu, Y., Turck, F., Li, L., Lu, Z., & Liu, L. (2021). Photoperiod-responsive changes in chromatin accessibility in phloem companion and epidermis cells of Arabidopsis leaves. Plant Cell, 33(3), 475–491. doi:10.1093/plcell/koaa043

Uhrig, R. G., Echevarria-Zomeno, S., Schlapfer, P., Grossmann, J., Roschitzki, B., Koerber, N., Fiorani, F., & Gruissem, W. (2021). Diurnal dynamics of the Arabidopsis rosette proteome and phosphoproteome. Plant Cell Environ, 44(3), 821–841. doi:10.1111/pce.13969

Uhrig, R. G., Schlapfer, P., Roschitzki, B., Hirsch-Hoffmann, M., & Gruissem, W. (2019). Diurnal changes in concerted plant protein phosphorylation and acetylation in Arabidopsis organs and seedlings. Plant J, 99(1), 176–194. doi:10.1111/tpj.14315

Veluchamy, A., Jegu, T., Ariel, F., Latrasse, D., Mariappan, K. G., Kim, S. K., Crespi, M., Hirt, H., Bergounioux, C., Raynaud, C., & Benhamed, M. (2016). LHP1 Regulates H3K27me3 Spreading and Shapes the Three-Dimensional Conformation of the Arabidopsis Genome. PLoS One, 11(7), e0158936. doi:10.1371/journal.pone.0158936

Venkat, A., & Muneer, S. (2022). Role of Circadian Rhythms in Major Plant Metabolic and Signaling Pathways. Front Plant Sci, 13, 836244. doi:10.3389/fpls.2022.836244

Wang, F., Han, T., & Jeffrey Chen, Z. (2024). Circadian and photoperiodic regulation of the vegetative to reproductive transition in plants. Commun Biol, 7(1), 579. doi:10.1038/s42003-024-06275-6

Wierzbicki, A. T., & Jerzmanowski, A. (2005). Suppression of histone H1 genes in Arabidopsis results in heritable developmental defects and stochastic changes in DNA methylation. Genetics, 169(2), 997–1008. doi:10.1534/genetics.104.031997

Yang, Y., Li, Y., Sancar, A., & Oztas, O. (2020). The circadian clock shapes the Arabidopsis transcriptome by regulating alternative splicing and alternative polyadenylation. J Biol Chem, 295(22), 7608–7619. doi:10.1074/jbc.RA120.013513

Zemach, A., Kim, M. Y., Hsieh, P. H., Coleman-Derr, D., Eshed-Williams, L., Thao, K., Harmer, S. L., & Zilberman, D. (2013). The Arabidopsis nucleosome remodeler DDM1 allows DNA methyltransferases to access H1-containing heterochromatin. Cell, 153(1), 193–205. doi:10.1016/j.cell.2013.02.033

