## supplementary figures and legend for "Linker Histones Enhance Robustness in Diurnal Transcription Dynamics"

Rutowicz et al.,

**Supplemental Information**

- Supplemental Figures 1-3
- Supplemental File 1 and Supplementary Figure S4 (description of the *3h1^crispr^* line)

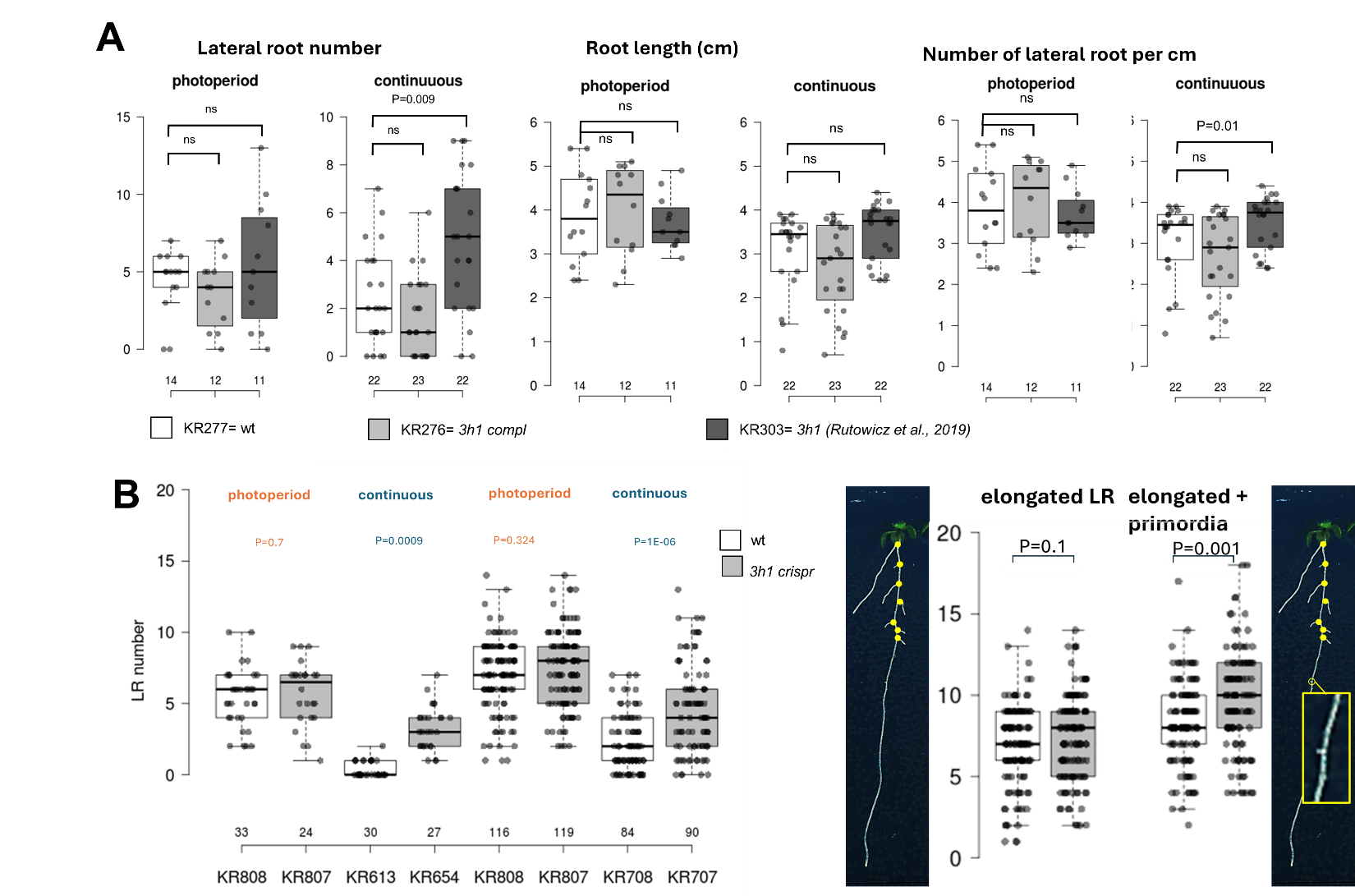

**Supplementary Figure S1A-B). The *3h1* mutant responds to the photoperiod at the phenotypic and transcriptomic level.**

(**A**) Lateral root number, root length, and lateral roots per cm per seedlings (dot) for 7-days-old wild-type seedlings (KR277, *3h1* segregant), *3h1* mutant (KR276), and *3h1* mutant (T-DNA allele) complemented with *pH1.1::H1.1-GFP* and *pH1.2::H1.2-CFP* (KR303) (Rutowicz et al., 2019), grown under long-day photoperiod or continuous light conditions as indicated. (**B**) Left graph: replicate measurements of lateral root scoring in 12 days old seedlings grown under the photoperiod or 7 days old seedlings grown under continuous light as indicated. White, wt seedlings segregants from the *3h1crispr* mutant; grey, *3h1 crispr* mutant seedlings. Right graph and pictures: the *3h1 crispr* mutant shows no significant difference in the number of elongated lateral roots (left image, yellow dots) with the wild-type whereas when young lateral root primordia are included in the scoring (right image, inlet), a slight difference is observed. Scoring was done on the same seedlings grown under a long-day photoperiod, n(wt)=116, n(3h1)=119. (A-B) Numbers above the X-axis indicate the number of seedlings analyzed (dots on the graph). P-values were determined by Kruskal-Wallis test with Bonferroni correction. See also **Supplemental Data 1**

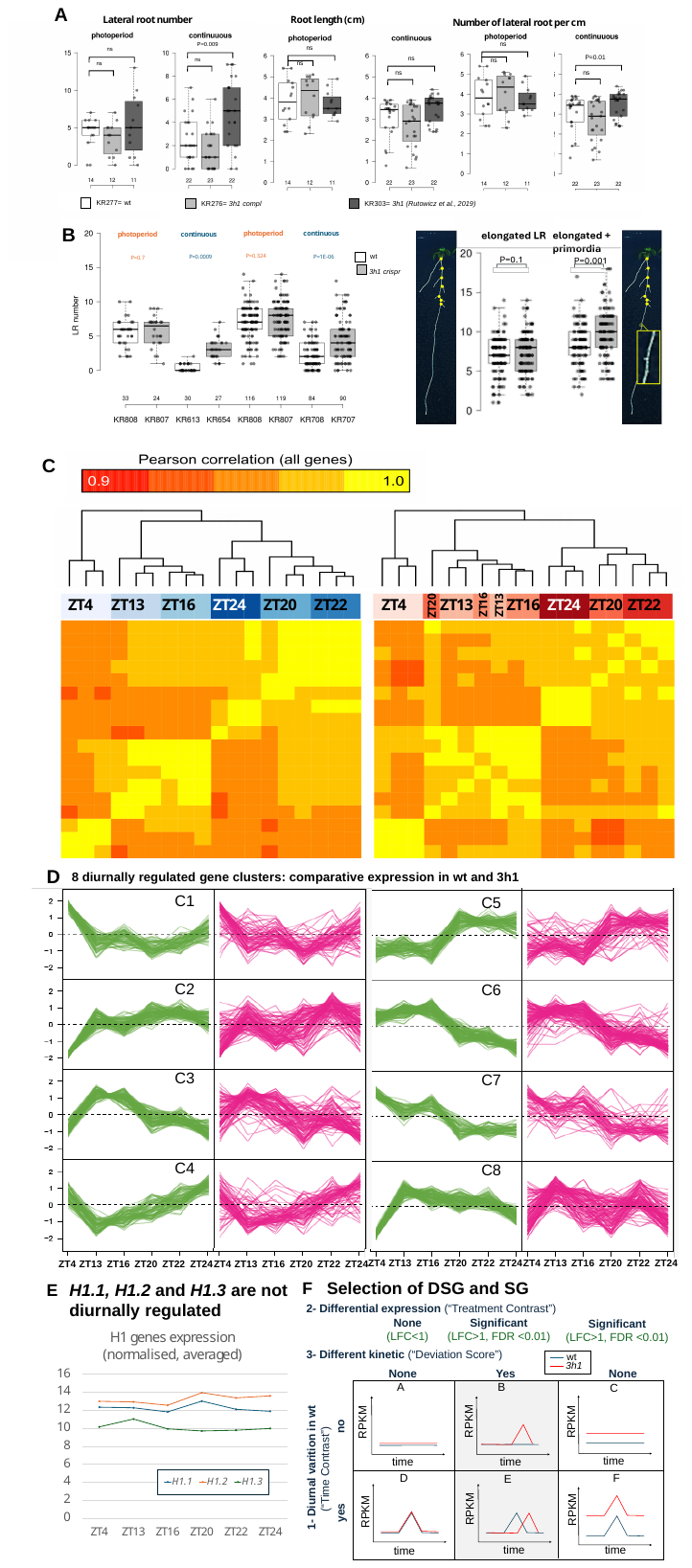

**Supplementary Figure S1(C-E). The *3h1* mutant responds to the photoperiod at the phenotypic and transcriptomic level.**

**Supplementary Figure S1(C-E). The *3h1* mutant responds to the photoperiod at the phenotypic and transcriptomic level.**

(**C**) Correlation analysis of transcriptome profiles from 19-day-old wild-type (blue) and *3h1* mutant (red) seedlings in three biological replicates at six time points (ZT4,13,16,20,22,24). Pearson correlation values range from 0.9 to 1. (**D**) Diurnal kinetics of differentially expressed genes grouped into eight clusters (C1-C8) in wild-type (green traces) and comparison to mutant samples (magenta traces). (**E**) Normalized, averaged expression from the 3 replicates in wt samples for the three H1 encoding genes, *H1.1(AT1G06760), H1.2(AT2G30620), H1.3(AT2G18050)* (**F**) Schematic of gene expression changes in wild-type (blue traces) and *3h1* mutant (red traces) across the photoperiod. The identification strategy for De-Synchronized Genes (DSG) and Synchronized Genes (SG) involves selecting genes with diurnal variation in wild-type, then identifying genes with differential expression between wild-type and mutant at any time point, followed by calculation of a deviation score to rank genes by their overall kinetic difference. The top and bottom 5% genes are classified as DSG (De-Synchronized) or SG (Synchronized), as shown in Figure 1. See also **Supplemental Data 1**

**Supplementary Figure S2. Characteristics of DSG vs SG groups**

(**A**) Left: Expression levels of DSG according to the DIURNAL database (Mockler et al., 2007); Right: One third of DSG are light-responsive (Schivre et al., 2025).(**B**) Normalised expression levels of DSG (n=197) and SG (n=773), the consecutive time-points t1-t6 correspond to ZT4, ZT13, ZT16, ZT20, ZT22, ZT24 (**C**) Metaplot enrichment analysis for SG and DSG, 2kb upstream and 2kb downstream gene start (GS) and gene end (GE), respectively, for H3K27me3 at day start (Baerenfaller et al., 2016) and H1 at an unspecified time point (Bourguet et al., 2021), and Venn-diagramm showing the overlap between DSG and hypo- or hyper-H3K27 methylated loci in the *2h1* mutant (Teano et al., 2023), (**D**) Metaplot enrichment analysis for SG and DSG of H3K4me3 across different time points (Song et al., 2019). (**E**) DNA methylation in each of the three contexts as indicated, at DSG and SG loci: enrichment levels in wt seedlings (upper panel) and differential enrichment between wt and *3h1* mutant seedlings (lower panel), Data from (Rutowicz et al., 2015), P values from a 2-side t-test. Values on the graph indicate the median. (**F**) Normalised expression levels (this study) of 17 NAC transcription factors which motives are most strongly enriched among DSG’s, showing absence of significant diurnal variations. See also **Supplemental Data 2**

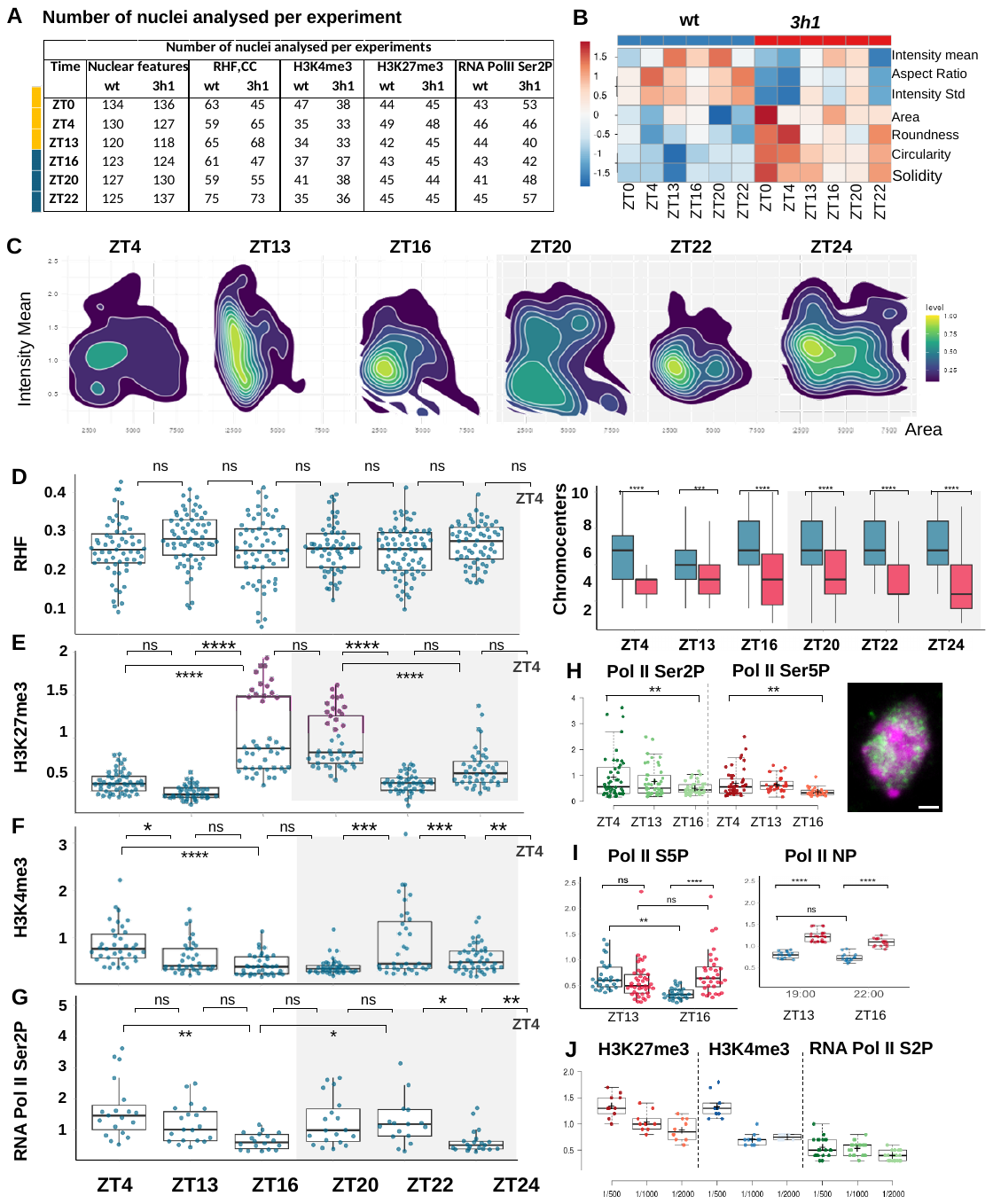

**Supplementary Figure S3. Quantification of diurnal chromatin changes in wt and *3h1***

Nuclei were isolated from seedlings harvested at the time points indicated, fixed on microscopic slides, imaged at high-throughput following immunostaining and DNA counterstaining, and batch-segmented (nucleus mask and chromocenters) using Nucl.Eye.D (Johann To Berens et al., 2022) for quantitative analyses. (**A**) Number of nuclei analysed in each experiment, comprising two biological replicate extractions, per time point and genotype. (**B**) Relative changes in nuclear features as listed on the right, along day and night, for each genotype. Heatmap generated by ClustVis (Metsalu & Vilo, 2015). Rows are centered and unit variance scaling is applied to rows. Both rows and columns are clustered using correlation distance and average linkage. (**C**) Changes in nuclear size (Area) and chromatin density (DNA Intensity mean) at different time points (ZT, X-axis) during the photoperiod, in wt seedling nuclei. Scatter plots with density contours generated by DataViz (Randall et al., 2022). (**D**) Quantification of the relative heterochromatin fraction (RHF) and number of chromocenters (CC), (**E-G**) Relative levels of H3K4me3, H3K27me3 and RNA Pol II Ser2P in wt, expressed as the ratio of antibody signal to DNA signal per nucleus, along the same time-points. (**H-I**) Quantification of RNA Pol II Ser2P, Ser5P or NP (non-phosphorylated) at selected time-points (ZT) in wt (H) or wt (blue) vs *3h1* (red) (I). (**J**) Comparison of immunostaining signal - relative to DNA intensity (Y-axis) for different dilutions of the secondary antibody as indicated on the X-axis. The differences between time points were evaluated using an ANOVA with Tukey HSD. ns, not significant. *, P<0.05. **, P<0.01. ***, P<0.001. ****, P<0.0001 See also **Supplemental Data 3**.

**Supplemental File 1 . Description of the *3h1^crispr^* mutant generated for the study**

Induction of targeted deletions, Arabidopsis Col_0 transformation and selection of mutant lines was done essentially as described (Bieluszewski et al., 2022). Guide RNAs against all three *H1.1, H1.2* and *H1.3* genes were co-expressed with Cas9 from a single genetic construct created by insertion of 6 gRNA expression cassettes into the pFGC-I2Cas9 binary vector (Addgene # 173158). The following sites were targeted:

H1.1 gRNA 1: GAGAACGCTGCTACGATCGA(AGG)

H1.1 gRNA 2: GTCTTTCGTCCTTTCGCTGC(AGG)

H1.2 gRNA 1: GTTCCAACGACTGTTGACTC(AGG)

H1.2 gRNA 2: GTTGGAGCAGCAGCTTTCAC(AGG)

H1.3 gRNA 1: GATGATAAAAGAGGCTTTGA(TGG)

H1.3 gRNA 2: AGTGTTTTACGGAAACTCTC(TGG)

PAM sequences in parentheses.

PCR-based screening of candidate mutant lines and combination of interesting alleles by crossing allowed to identify single and double mutants but no triple mutants. Thegenerate the new 3h1crispr line, which was subsequently was sgenerated by crossing a double mutant and a single mutant in the complementary allele and cleaned from the CAS9-expressing vector through two successive backcrossing. Individual *h1.1^crispr^*, *h1.2^crispr^* and *h1.3^crispr^* alleles were obtained by backcrossing.

The selected *h1.1^crispr^* mutant allele contains a 61 bp deletion in the first exon; the selected *h1.2^crispr^* allele contains a 54bp deletion and a 1bp insertion (leading to an effective 53 bp deletion) in the first exon; the selected *h1.3^crispr^* allele contains a 35bp deletion followed by a 2bp deletion (leading to an effective 37bp deletion). All mutations cause frameshifts in the coding region (**Supplementary Figure SS4A, B**).

Multiplex PCR combining primer pairs to genotype all three deletions was used for selection. A standard PCR with a primer pair specific to the CAS9-expressing vector was also used for selection. The table below show the set of primer used, expected product length in wild-type (wt) or mutant (*3h1*) plants. A typical gel is shown **Supplementary Figure SS4C**. The multiplex PCR-based genotyping assay uses 34 cycles of 30” denaturation at 94°C, 30” annealing at Tm=60°C and 80” elongation at 72°C)

| **Name** | **Sequence (5’-3’)** | **Product length (wt/3h1)** | **Primer length** | **Tm** | **GC%** |
| --- | --- | --- | --- | --- | --- |
| H1.1crF3 | acagttctcttcttcggagcgg | 169 / 108 | 22 | 62.28 | 54.55 |
| H1.1crR3 | aggtggaaatagagaacgctgct | 169 / 108 | 23 | 61.95 | 47.83 |
| H1.2crF1 | catcacaaaattctccgacgca | 287 / 234 | 22 | 59.84 | 45.45 |
| H1.2crR1 | tcgaatattgacctcttcataggtagg | 287 / 234 | 27 | 59.82 | 40.74 |
| H1.3crF1 | accaccactcatcctccatact | 394 / 357 | 22 | 60.29 | 50.00 |
| H1.3crR1 | gccttgtcgaagaagacctagt | 394 / 357 | 22 | 59.77 | 50.00 |
| InsCr3h1-F1 | gatggactataaggaccacgacg | 102 | 23 | 60.30 | 52.17 |
| InsCr3h1-R1 | ataccgaccttccgcttcttctt | 102 | 23 | 61.69 | 47.83 |

Known mutant alleles generated by insertional mutagenesis (T-DNA) were reported previously with occasionally some inconsistencies regarding their name and numbering. We propose to follow the nomenclature and give the new alleles a number corresponding to their rank in the list of alleles described by published studies.

- ***h1.1 alleles***:
  - *SALK_128430C (N628430) described as* ***h1.1-1*** *in (Han et al., 2022; Rutowicz et al., 2015)*
  - *h1.1^crispr^ described here as* ***h1.1-2***
- ***h1.2 alleles****:*
- *GK-116E08 described as* ***h1.2*** *in (Rutowicz et al., 2015)*
- SALK_002142 *described as* ***h1.2-1*** *in (Han et al., 2022)*
- CS438975 described as ***h1.2-2*** *in (Han et al., 2022)*
- GABI_406H11 described as ***h1.2*** in (Bourguet et al., 2021)
- *h1.2^crispr^ described here as* ***h1.2-5***
- ***h1.3 alleles***
- SALK_025209 *described as* ***h1.3-1*** *in (Han et al., 2022)*
- GT18298 described as *h1.3 (Ler)* in (Rutowicz et al., 2015)
- *h1.3^crispr^ described here as* ***h1.1-3***

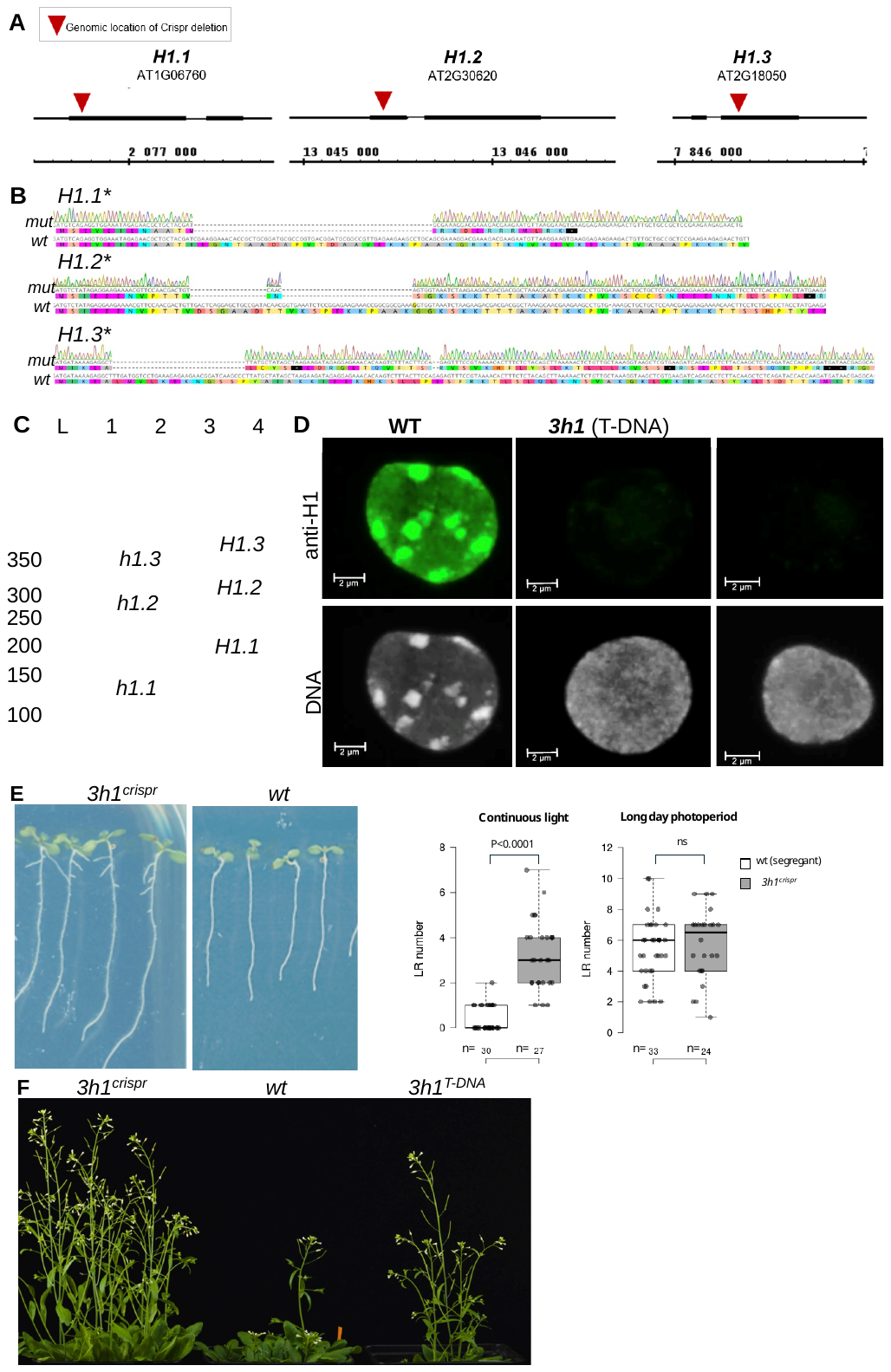

**Supplementary Figure SS4 - The 3h1^crispr^ mutant recapitulates known phenotypes of the 3h1^T-DNA^ mutant**

(**A-B**) The newly generated 3h1^crispr^ mutant line carries small deletions in 61 bp, 53 bp and 37 bp deletions in the first exon of *H1.1, H1.2* and second exon of *H1.3* respectively. (**C**) Multiplex genotyping assay on leaf DNA extract using primer conditions as detailed in the text. L, 50bp Ladder Thermo Scientific GeneRuler 50 bp DNA Ladder); 1-2, two 3h1^crispr^ mutant samples; 3-4, two wild-type samples. Note the smaller size of all three amplicons in the mutant due to the deletions in *H1.1, H1.2 and H1.3*, respectively. (**D**) Isolated leaf nuclei were immunostained using a rabbit anti-H1 (AS11-1801, Agrisera, Sweden) antibody (green) as described in (She et al., 2013) and counterstained with Propidium Iodide (grey) before confocal imaging. Images show 3D max projections. Both the T-DNA insertion line (Rutowicz et al., 2019) and newly created 3h1^crispr^ line used for this study show decondensed heterochromatin and undetectable H1 immunosignal. (**E-F**) The newly created 3h1^crispr^ line used for this study recapitulates the lateral root and early flowering phenotypes as reported previously (Rutowicz et al., 2019). The number of lateral roots was scored for 7 and 12 days old seedlings grown under continuous light or a long day photoperiod, respectively (n, number of seedling scored; P values from a Mann-Whitney U test).

**References**

Baerenfaller, K., Shu, H., Hirsch-Hoffmann, M., Futterer, J., Opitz, L., Rehrauer, H., Hennig, L., & Gruissem, W. (2016). Diurnal changes in the histone H3 signature H3K9ac|H3K27ac|H3S28p are associated with diurnal gene expression in Arabidopsis. *Plant Cell Environ, 39*(11), 2557-2569. doi:10.1111/pce.12811

Bieluszewski, T., Sura, W., Dziegielewski, W., Bieluszewska, A., Lachance, C., Kabza, M., Szymanska-Lejman, M., Abram, M., Wlodzimierz, P., De Winne, N., De Jaeger, G., Sadowski, J., Cote, J., & Ziolkowski, P. A. (2022). NuA4 and H2A.Z control environmental responses and autotrophic growth in Arabidopsis. *Nat Commun, 13*(1), 277. doi:10.1038/s41467-021-27882-5

Bourguet, P., Picard, C. L., Yelagandula, R., Pelissier, T., Lorkovic, Z. J., Feng, S., Pouch-Pelissier, M. N., Schmucker, A., Jacobsen, S. E., Berger, F., & Mathieu, O. (2021). The histone variant H2A.W and linker histone H1 co-regulate heterochromatin accessibility and DNA methylation. *Nat Commun, 12*(1), 2683. doi:10.1038/s41467-021-22993-5

Johann To Berens, P., Schivre, G., Theune, M., Peter, J., Sall, S. O., Mutterer, J., Barneche, F., Bourbousse, C., & Molinier, J. (2022). Advanced Image Analysis Methods for Automated Segmentation of Subnuclear Chromatin Domains. *Epigenomes, 6*(4). doi:10.3390/epigenomes6040034

Metsalu, T., & Vilo, J. (2015). ClustVis: a web tool for visualizing clustering of multivariate data using Principal Component Analysis and heatmap. *Nucleic Acids Res, 43*(W1), W566-570. doi:10.1093/nar/gkv468

Mockler, T. C., Michael, T. P., Priest, H. D., Shen, R., Sullivan, C. M., Givan, S. A., McEntee, C., Kay, S. A., & Chory, J. (2007). The DIURNAL project: DIURNAL and circadian expression profiling, model-based pattern matching, and promoter analysis. *Cold Spring Harb Symp Quant Biol, 72*, 353-363. doi:10.1101/sqb.2007.72.006

Randall, R. S., Jourdain, C., Nowicka, A., Kaduchova, K., Kubova, M., Ayoub, M. A., Schubert, V., Tatout, C., Colas, I., Kalyanikrishna, Desset, S., Mermet, S., Boulaflous-Stevens, A., Kubalova, I., Mandakova, T., Heckmann, S., Lysak, M. A., Panatta, M., Santoro, R., Schubert, D., Pecinka, A., Routh, D., & Baroux, C. (2022). Image analysis workflows to reveal the spatial organization of cell nuclei and chromosomes. *Nucleus, 13*(1), 277-299. doi:10.1080/19491034.2022.2144013

Rutowicz, K., Puzio, M., Halibart-Puzio, J., Lirski, M., Kotlinski, M., Kroten, M. A., Knizewski, L., Lange, B., Muszewska, A., Sniegowska-Swierk, K., Koscielniak, J., Iwanicka-Nowicka, R., Buza, K., Janowiak, F., Zmuda, K., Joesaar, I., Laskowska-Kaszub, K., Fogtman, A., Kollist, H., Zielenkiewicz, P., Tiuryn, J., Siedlecki, P., Swiezewski, S., Ginalski, K., Koblowska, M., Archacki, R., Wilczynski, B., Rapacz, M., & Jerzmanowski, A. (2015). A Specialized Histone H1 Variant Is Required for Adaptive Responses to Complex Abiotic Stress and Related DNA Methylation in Arabidopsis. *Plant Physiol, 169*(3), 2080-2101. doi:10.1104/pp.15.00493

Schivre, G., Wolff, L., Mirasole, F. M., Armanet, E., Davidson, M. L. H., Vidal, A., Cuménal, D., Dumont, M., Bourge, M., Baroux, C., Bourbousse, C., & Barneche, F. (2025). Genome-scale transcriptome augmentation during *Arabidopsis thaliana* photomorphogenesis. *bioRxiv*, 2025.2001.2030.635720. doi:10.1101/2025.01.30.635720

Song, Q., Huang, T. Y., Yu, H. H., Ando, A., Mas, P., Ha, M., & Chen, Z. J. (2019). Diurnal regulation of SDG2 and JMJ14 by circadian clock oscillators orchestrates histone modification rhythms in Arabidopsis. *Genome Biol, 20*(1), 170. doi:10.1186/s13059-019-1777-1

Teano, G., Concia, L., Wolff, L., Carron, L., Biocanin, I., Adamusova, K., Fojtova, M., Bourge, M., Kramdi, A., Colot, V., Grossniklaus, U., Bowler, C., Baroux, C., Carbone, A., Probst, A. V., Schrumpfova, P. P., Fajkus, J., Amiard, S., Grob, S., Bourbousse, C., & Barneche, F. (2023). Histone H1 protects telomeric repeats from H3K27me3 invasion in Arabidopsis. *Cell Rep, 42*(8), 112894. doi:10.1016/j.celrep.2023.112894
